# A laser capture microdissection transcriptome of *M. truncatula* roots responding to rhizobia reveals spatiotemporal tissue expression patterns of genes involved in nodule signaling and organogenesis

**DOI:** 10.1101/2023.03.14.532650

**Authors:** Elise Schnabel, Jacklyn Thomas, Rabia El-Hawaz, Yueyao Gao, William Poehlman, Suchitra Chavan, Asher Pasha, Eddi Esteban, Nicholas Provart, F. Alex Feltus, Julia Frugoli

## Abstract

We report a public resource for examining the spatiotemporal RNA expression of 54,893 *M. truncatula* genes during the first 72 hours of response to rhizobial inoculation. Using a methodology that allows synchronous inoculation and growth of over 100 plants in a single media container, we harvested the same segment of each root responding to rhizobia in the initial inoculation over a time course, collected individual tissues from these segments with laser capture microdissection, and created and sequenced RNA libraries generated from these tissues. We demonstrate the utility of the resource by examining the expression patterns of a set of genes induced very early in nodule signaling, as well as two gene families (CLE peptides and nodule specific PLAT-domain proteins) and show that despite similar whole root expression patterns, there are tissue differences in expression between the genes. Using a rhizobial response data set generated from transcriptomics on intact root segments, we also examined differential temporal expression patterns and determined that, after nodule tissue, the epidermis and cortical cells contained the most temporally patterned genes. We circumscribed gene lists for each time and tissue examined and developed an expression pattern visualization tool. Finally, we explored transcriptomic differences between the inner cortical cells that become nodules and those that do not, confirming that the expression of ACC synthases distinguishes inner cortical cells that become nodules and provide and describe potential downstream genes involved in early nodule cell division.

## Introduction

Legume plants establish a symbiotic relationship with rhizobia to access fixed atmospheric nitrogen when available soil nitrate is low, allowing legumes to be grown without added nitrogen fertilizer. The signal transduction events in the legume-rhizobial symbiosis not only involve organisms of two different kingdoms (bacteria and plants) but the communication required to establish the symbiosis occurs between cells layers in tissues, between organs in the plant, and across time, from the induction of the first chemical responses within 6-12 hours to the establishment of nitrogen fixation in the nodules 10 days after inoculation (reviewed in Roy et al., 2020, Yang, et al. 2022). Each organism influences the development of the other, resulting in differentiated bacteria living inside the cells of a plant organ (a nodule) that allows the bacteria to reproduce and fix nitrogen by providing carbon skeletons and a pocket of low oxygen tension.

Early symbiotic signaling starts with the initial signal transduction in root hair cells. Briefly summarized from (Oldroyd, 2013; Yang et al., 2022), rhizobia produce species-specific lipo-chitin oligosaccharides called Nod factors in response to flavonoids secreted into the soil by legumes, and this initiates the symbiotic interaction. In response to the plant flavonoids secreted into the soil, the bacteria chemotax to the root and attach to the root hairs, where they begin to produce Nod factor. Depending on the sugar decorations and saturation of the N acyl groups of Nod factor, the plant detects Nod factor as coming from a compatible species of rhizobia with a LysM receptor-like kinase complex. SYMREM (symbiotic remorin), FLOT2 and FLOT4 (flotillins) are predicted to bring the Nod factor receptors into lipid rafts (Haney and Long, 2010; Lefebvre et al., 2010). In *M. truncatula*, the complex that detects Nod factor contains several receptors including NFP and LYK3 with non-functional and functional kinase domains, respectively, that detect Nod factor, and DMI2 which signals through 3-hydroxy-3methylglutaryl-CoA reductase and other proteins to the nucleus (Oldroyd, 2013; Venkateshwaran et al., 2015)

Within minutes of Nod factor detection, calcium oscillations begin in the nucleus (Ehrhardt et al., 1996), where the nuclear channel DMI1 and nuclear porins have been genetically shown to play a role in this signal transduction event (Capoen et al., 2011). The calcium oscillation connects the receptor signaling at the plasma membrane to the downstream signaling events through the calcium-calmodulin dependent kinase DMI3 (Levy et al., 2004). DMI3 phosphorylates IPD3 which is essential for microbial colonization (Horvath et al., 2011). The GRAS family transcription factors NSP1 and NSP2 are stimulated to form a heterocomplex that activates nodulation transcription factors (NIN and ERN1) as well as other nod factor inducible genes such as *ENOD11*, and DELLA, which is able to bridge a protein complex containing IPD3 and NSP2 (Jin et al., 2016).

The majority of what is known about early symbiotic signaling involves events which occur within a few hours of infection in the elongation zone of the root. Root hairs in cells that have ceased to elongate do not respond to rhizobia (Gage, 2004). Upon attachment of the rhizobia to the root hair tips, the root hairs curl tightly and entrap the bacteria in the curl. The plant cell forms a new structure, a tubular infection thread, through which the bacteria enter the plant through cell division. At the same time as infection thread formation, a subset of the inner cortical cells next to the xylem poles are mitotically activated; these cells will eventually form the nodule primordia. The nature of the signal that reactivates the cell cycle is unclear, although it is likely to be a component of the cell cycle (Murray, 2016) and *ENOD40* is required (Charon et al., 1997). However, *in situ* analysis of ACC synthase activity in wild type plants and examination of ethylene insensitive *M. truncatula* mutants (*sickle*, carrying a disruption of the *EIN2* ortholog) strongly suggest that this positional information is related to ethylene levels in the cortical cells between the xylem poles (Heidstra et al., 1997; Penmetsa and Cook, 1997; Penmetsa et al., 2008).

Within the next 24 hours, the infection threads cross the outer cortical cells and begin to branch. The outer cortical cells undergo rearrangements reminiscent of phragmoplast formation that allow the infection threads to pass through the cells (Brewin, 1991). Only a subset of initiated infection threads will persist into the inner cortex; the majority will arrest in the outer cortex due to an unknown mechanism (Gage, 2004). Meanwhile, the inner cortical cells continue to divide, and the concentrations of both cytokinin and auxin change in both directions in the cortex and endodermis/pericycle area (reviewed in Lin et al., 2020). Whole roots measurements show an increase in cytokinin levels (reviewed in (Gamas et al., 2017) and a reduction of auxin transport from the shoot to the root (van Noorden et al., 2006), but this has not been examined in detail at the level of individual cells, except to note that Nod factor affects polar auxin transport only when applied to the elongation zone of the root, where the nodules will form (Suzaki et al., 2012).

By 48 hours after inoculation the nodule primordia have begun to organize into a meristematic region flanked by a region of cells that have ceased dividing. In *M. truncatula*, the genes encoding CLE12 and CLE13 peptides are expressed in the meristematic area (Mortier et al., 2010) and are involved in autoregulation, sending a nodulation signal to the shoot (Okamoto et al., 2013). Local changes in auxin transport in the vascular bundle occur where nodules are forming (Mathesius et al., 1998; Liu et al., 2018) similar to what happens when lateral roots initiate, as well as expression of a subset of auxin transporter genes (*PIN* genes) required for nodule and lateral root development (Huo et al., 2006). *LATERAL ORGAN BOUNDARY* (*LBD*) and *STYLISH*-like (*STY-l*) genes also promote cell proliferation leading to the nodule (Schiessl et al., 2019).

At 72 hours after inoculation, *M. truncatula* will halt the initiation of additional nodule primordia in the elongation zone, presumably in response to an autoregulatory signal (Gage, 2004; Kassaw and Frugoli, 2012). Under controlled conditions this results in a fixed number of nodules in a small area of the root corresponding to the maturation zone at the time of inoculation (Bhuvaneswari et al., 1980). Bacteria in infection threads that have entered the outer cortex stop dividing and the threads degrade, auxin transport resumes a normal pattern, and the successful infection threads will begin to release bacteria into symbiosomes in the cells that have ceased to divide behind the meristem (Gage, 2004). The developing nodule is still within the cortex and the vasculature that will feed it has not yet organized, but from this point on the development of the nodule seems to be controlled by signals from the meristem and signals from the bacteria within the symbiosomes.

For much of nodule development after the first few hours, the physical progression is known from microscopy observations, yet until very recently the underlying molecular signals were only known at a gross, whole root level. While gene regulation analyses have advanced our knowledge of the nodule initiation process (reviewed in Mergaert et al., 2020), and transcriptome profiling of whole roots at one or two ‘snapshot’ time points during nodule development has expanded knowledge of what genes are involved in initiating symbiosis, such experiments are unable to resolve the progression of events at the tissue level where many sub-organ gene expression signals are likely diluted by the complex root tissue mixture. For example, a subset of successful infection threads passes through the root epidermis and the outer cortical cells to the inner cortical cells. At a spatial level, the cell cycle has reactivated in inner cortical cells next to the xylem poles but not in inner cortical cells next to the phloem, and at a temporal level the cell division begins in these cells in anticipation of the arrival of the thread, one branch of which will penetrate to the nodule (reviewed in Gage, 2004)). The vasculature is separated from the inner cortical cells by the pericycle and endodermis, so spatial signals must be selectively passed across the pericycle and endodermis and be communicated to the infection thread proceeding across the cortex.

To understand the signaling occurring between these tissues in space and time in a coherent manner, we analyzed the transcriptome of the individual tissues at specific points in time, discovering that neighboring tissues have different transcriptomes. Using Laser Capture Microdissection (LCM) of nodulating roots over five early time points, we were able to create high resolution gene expression profiles to further elucidate tissues and factors involved in early nodulation signaling. Two previous experiments using LCM in *M. truncatula* compared colonized and non-colonized cells undergoing arbuscular mycorrhizal infection, then used microarray hybridization to detect genes and published a catalog of differential expression (Gomez et al., 2009; Gaude et al., 2012). Two other groups published several analyses using LCM and RNASeq to look at both bacterial and plant gene expression in late nodulation (nodules 15 days post inoculation) (Limpens et al., 2013; Roux et al., 2014; Jardinaud et al., 2016).

Our study examines critical signal transduction time points of *early nodulation* signaling with the single tissue information of LCM and the resolution power of RNASeq. While a recent paper on single-cell RNASeq of nodulating *M truncatula* roots at a single time point of 48 hours post inoculation has added to our knowledge (Cervantes-Pérez et al., 2022), the work presented here takes a different but complementary approach. We also provide a public visualization interface (https://bar.utoronto.ca/eplant_medicago/) which allows queries of the database for the timing and tissue of all genes whose expression we could detect during early nodulation. We used the interface to present new and confirmatory information on the spatiotemporal patterning of known nodulation genes such as *NFH1, VPY, NSP1, NIN, ERN1*, and *ENOD11* as well as examining the variation in pattern among members of the *CLE* and *NPD* gene families. Dissecting the distinct patterning of rhizobial response by tissue revealed many patterns in the epidermis and cortical cells and allowed for the identification of genes underlying the observation that nodules form in the inner cortex next to the xylem poles versus in the inner cortical space in between the poles.

## Results

### Physical progression of nodule development during this time course

To capture signaling events in early nodule development, we developed a harvesting procedure described in (Schnabel et al., 2023). The procedure involved simultaneous inoculation of all plants in an aeroponic system (Cai, et al. 2023) and harvest of the developing nodule zone tracked via root growth (summarized in Supplemental Figure 1). In our experience, nodulation progresses more uniformly and rapidly in this system than on plates or in pots, because the apparatus sprays the aerosolized solution of rhizobia on all plants simultaneously. Cross sections of the root segments harvested for LCM at the various timepoints (Figure 1 A-E) showed developmental progress of the nodulation response at the times of collection. The 0-hour post inoculation (hpi) time point under all conditions is comparable between inoculated and uninoculated experiments (Figure 1A). Twelve hours post inoculation, some cellular response (occasional cell divisions) was observed (Figure 1B), and by 24 hpi the initial divisions of cells near the xylem pole that will become a nodule was clearly observed (arrow in Figure 1C). By 48 hpi nodule meristems formed (Figure 1D), and by 72 hpi nodules broke through the epidermal layer which is observable by the naked eye (Figure 1E).

**Figure 1.**
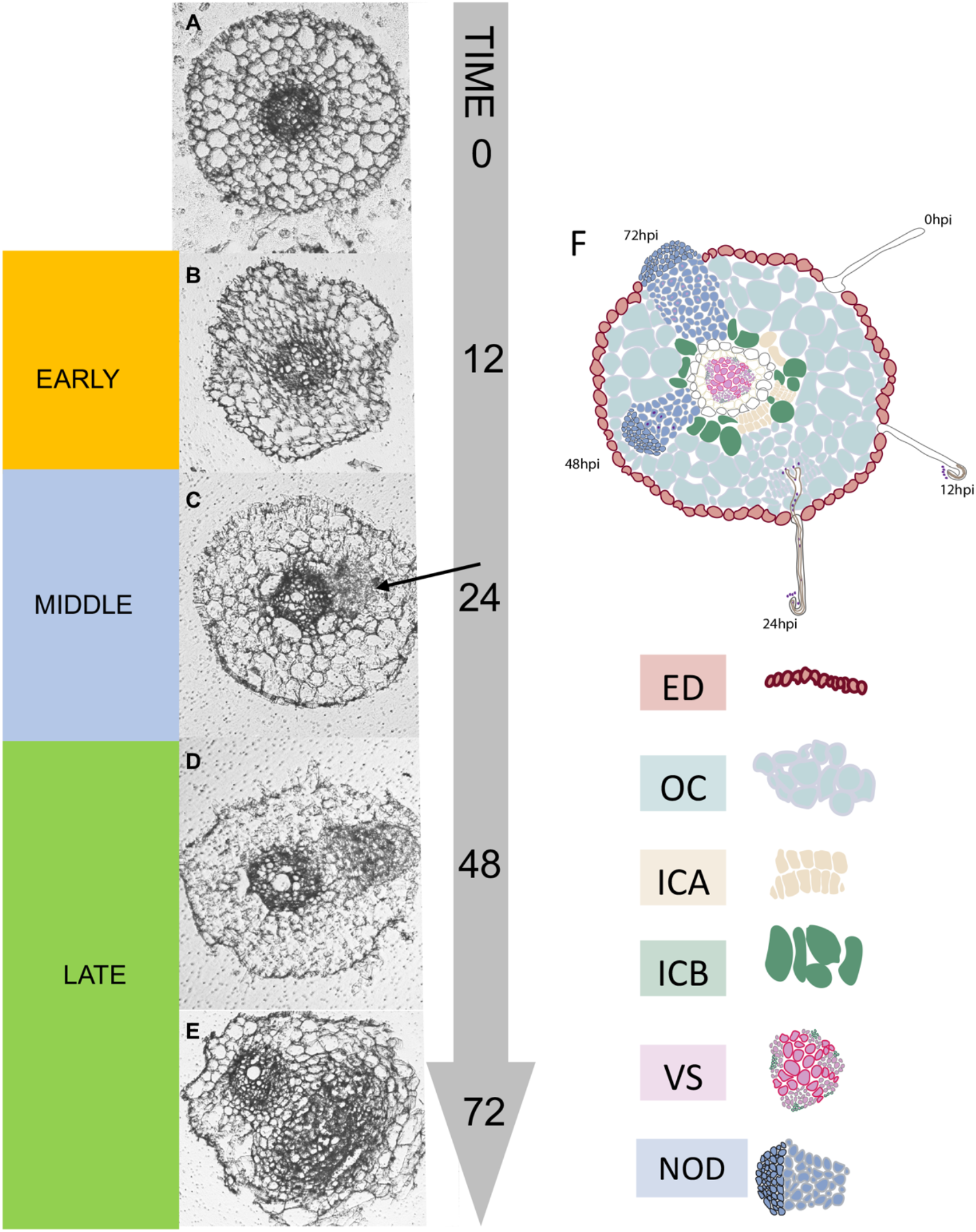
Stages of early nodule development and areas of tissue harvest. Images are micrographs of cross sections from root segments grown in an aeroponic system as described in methods and harvested at the indicated time points. Roots were fixed in and cut as in Chavan, et al. (2018). (A) uninoculated root (B) 12 hours post inoculation (hpi) (C) 24 hpi (arrow indicates early cell divisions) (D) 48 hpi (E) 72 hpi. Early (I), Middle (II), and Late (III) refer to categories for rhizobial response gene analysis (see Results). (F) Schematic of tissues captured for RNA at timepoints in this experiment. All tissue types were captured at each time point, except for NOD which was captured only at 48 and 72 hours post inoculation. Abbreviations used throughout the manuscript are ED=epidermal cells, OC=outer cortical cells, ICA= inner cortical cells across from xylem poles, ICB=inner cortical cells between xylem poles, VS=vasculature cells, NOD=cells forming a nodule.

In our analysis of rhizobial response in unfixed root segments reported later in results, we considered transcriptional changes that happen by 12 hpi as “Early,” 24 hpi as “Middle” and 48 to 72 hpi as “Late” to connect transcriptional events to stages of nodulation signaling and development. Figure 1F displays the locations of tissues captured at different time points from root sections. Note that “Nodule” tissue was only captured from 48- and 72-hpi root cross sections, and the remaining Epidermis (including root hairs) (ED), Vasculature (VS), Outer Cortical Cells (OC), and Inner Cortical Cells Between (ICB) and Across (ICA) from xylem poles at 48 and 72 hpi were captured from cross sections that did not include a nodule. This allowed us to treat the developing nodule separately from the rest of the root. In wild type plants, the cells across from the xylem poles become nodules, while those between the poles rarely become nodules; this pattern is correlated with ethylene signaling (Heidstra et al., 1997; Penmetsa and Cook, 1997; Penmetsa et al., 2008), but little else is known. Inner cortical cells were therefore separated into two categories based on spatial position to investigate nodule position signaling.

### Early nodulation genes vary in expression increases in different tissues

The data from this experiment (Supplemental Data Set 1) as well as data from (Schnabel et al., 2023) in which an analysis of gene expression changes in root segments during the same time period, were deposited at ePlant and display options created in ePlant (https://bar.utoronto.ca/eplant_medicago/) to allow analysis of the 4,893 individual genes in our dataset. Figure 2C is a representative visualization created by loading the example gene Medtr8g043970 (*ERN1*) into ePlant, selecting the tissue and experiment viewer (microscope icon), and then clicking on the Root Component view (root cross section) to generate the image. This public resource allows other investigators to examine gene expression in early nodulation timepoints in our data with a user-friendly interface (Waese et al., 2017).

**Figure 2.**
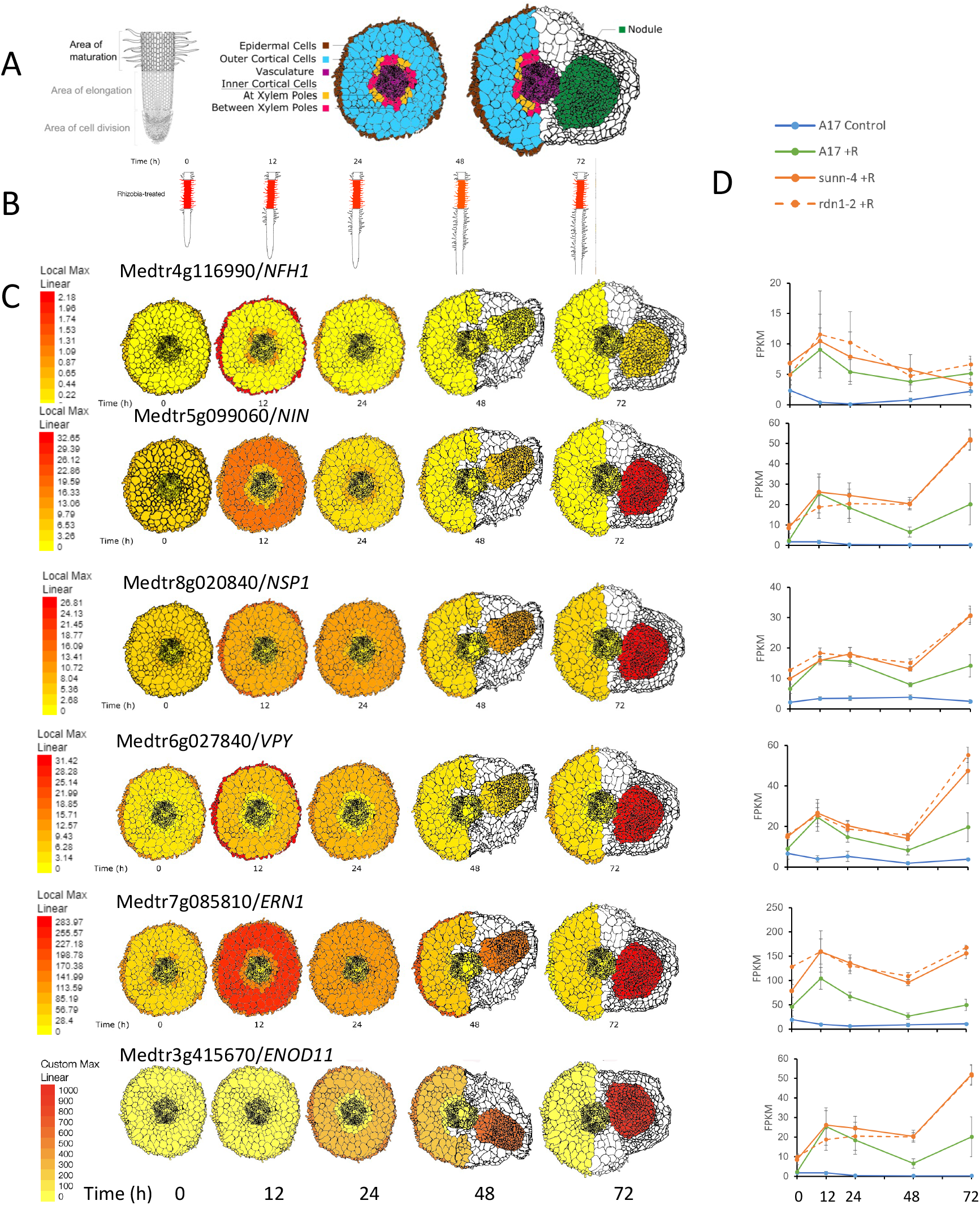
Induction of nodulation genes in tissue responding to Rhizobia. A) Diagram of the tissues sectioned, from the area of elongation at 0 hpi, and from the areas corresponding to the 0 hpi area at later time points. B) Corresponding area captured at each timepoint (also displayed in ePlant). C) Expression patterns of indicated genes at each timepoint by FPKM. Note the scale is calculated to a local maximum, indicated in red, which varies by the gene. D) Average expression, with error bars of the same genes in B in whole root segments under the same conditions (from data in Schnabel, et al 2023).

To confirm the quality of the data, we first examined tissue level expression of specific genes known to be involved in nodulation chosen from a recent review (Roy et al., 2020), using the ePlant interface. Figure 2A displays the area of tissue sampling (maturation zone at 0 hpi, and the same cells followed through 72 hpi) as well as which cells will be “painted” based on expression in that tissue. Each display paints a local maximum as red, with the gene expression values (FPKM) displayed on the scale bar to the left of each row. The user can change the display by changing the maximum to a manual entry (for instance to eliminate a single tissue such as nodules from overwhelming other expression differences as done for *ENOD11* below) or using a global maximum for the whole gene set to compare across genes in a single tissue. The genes in Figure 2B have different expression maximums (see scale bar on left), but the use of a local expression maximum for each gene aid tissue expression pattern comparisons versus comparing actual values. Figure 2C shows the expression of the same genes in 2B in whole root segments in inoculated and uninoculated wild type plants grown in the same system and protocol, as well as in the *sunn-4* and *rdn1-2* hypernodulation mutants. These graphs were generated from data in (Schnabel et al., 2023).

As expected, all the genes chosen for examination in Figure 2B showed an increase in expression at one point in the time course in Figure 2C that was not present in the absence of rhizobia (the blue line in Figure 2C). However, the specific tissue with an increase in expression varied by gene. For example, as expected given the central role of *NIN* in nodule signaling, *NIN* expression rose rapidly in the cortical cells, dropped by 24-48 hours, and then rose in the developing nodule at 72 hours. The *ERN1* gene showed the same pattern, although the expression level was an order of magnitude more than *NIN*. Meanwhile, *NFH1, NSP1*, and *VPY*, while they appear to have a similar expression pattern to *NIN* when the root segment traces in Figure 2C were examined, all displayed increased expression in epidermal cells but not cortical cells followed by the same rise in nodules as was observed for *NIN. ENOD11*, which is genetically downstream of *NSP1, NIN* and *ERN1*, exhibited a 100-fold higher expression in nodules than other tissues. For *ENOD11*, the display in Figure 2B was capped at half the expression at 72 hours in nodule tissue to help visualize the earlier tissue expression patterns, especially in the epidermis at 24 and 48 hpi. In all these cases, a similar temporal pattern in whole root segments was revealed by LCM as differential expression patterns in individual tissues.

### Gene family members display different tissue patterns

We then examined expression of two multi-gene families containing multiple members involved in nodulation to determine if family member-specific patterns of expression could be identified. The *CLE* gene family is a large family of peptide encoding genes dependent on processing for their biological activity, reviewed in (Roy and Müller, 2022). The *CLE* genes in Figure 3 have all been reported to affect nodule development or AM symbiosis (Roy and Müller, 2022) but *MtCLE53*, implicated in AM symbiosis, *peaked* early in wild type root segments responding to rhizobia and then declined, while the rest of the nodulation CLEs increased early and remain at this level or continued to rise (green lines in Figure 3B). When examined at the tissue level (Figure 3A), *MtCLE37, MtCLE12* and *MtCLE45* displayed maximum expression in nodules at later timepoints, while *MtCLE53* and *MtCLE35* showed maximum expression in the vasculature and inner cortex at earlier (12-24 hpi) timepoints. *MtCLE13* is an outlier, with maximum expression at 12 hpi in the cortex, followed by a secondary but slightly lower maximum at 48-72 hpi in nodules. A second gene family, nodule-specific PLAT domain genes, have functions in infection; *MtNPD1* is necessary for rhizobial accommodation and is found in the space surrounding the symbiosomes (Pislariu et al., 2019). *MtNPD2* and multiple mutants of *MtNDP3, MtNDP4* and *MtNDP5* all show nodulation phenotypes where the loss of all these genes eliminates nitrogen fixation (Trujillo et al., 2019). As expected, the expression of all *MtPLAT* genes began to increase as nodules developed between the 48 and 72 hpi time points (Figure 4B) in our root segment data. At a tissue level (Figure 4A) all nodule-specific *PLAT* domain genes showed maximum expression in nodules, but a close look at *MtNPD2* expression suggested a secondary maximum in outer cortical cells at 24 hpi, and *MtNDP3,4* and *5* all showed a secondary maximum in inner cortical cells at 24 hpi. Thus, the expression of *MtNDP* genes is nodule associated but not nodule specific.

**Figure 3.**
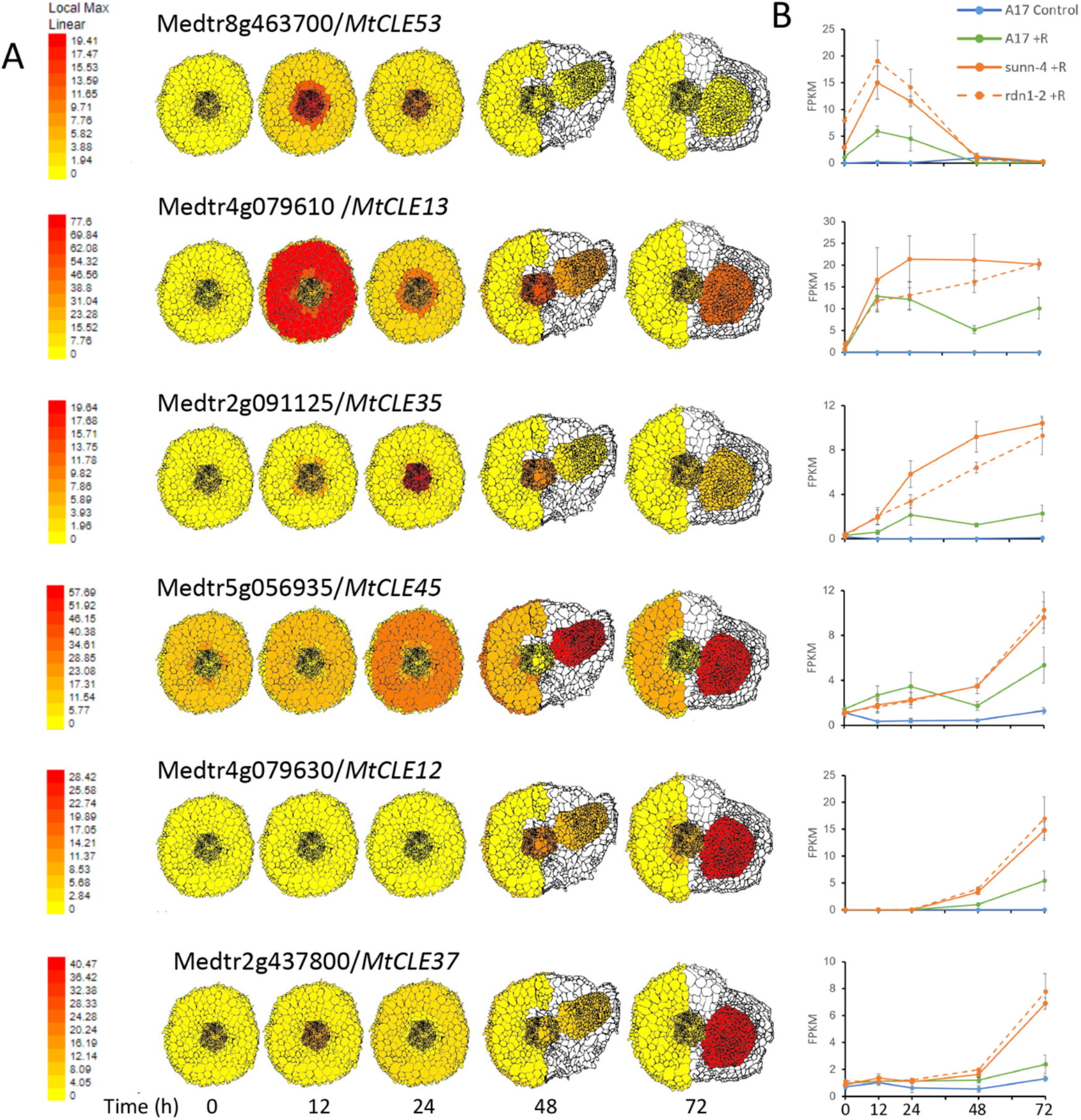
Induction of CLE Peptide Genes in Tissue Responding to Rhizobia. A) Expression patterns of indicated genes at each timepoint by FPKM. Note the scale is calculated to a local maximum, indicated in red, which varies by the gene. B) Average expression, with error bars of the same genes in B in whole root segments under the same conditions (from data in Schnabel, et al 2023).

**Figure 4.**
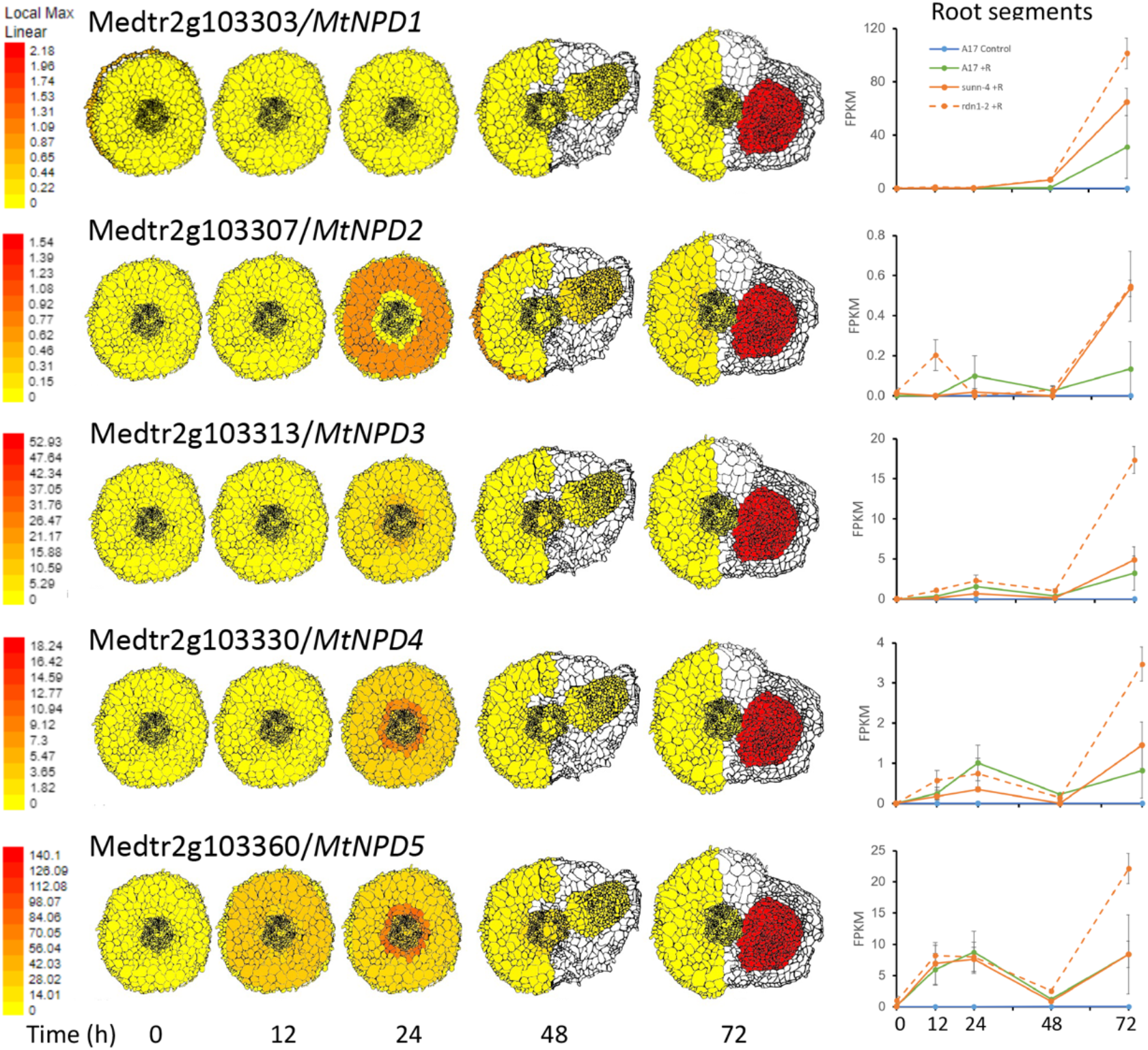
Induction of nodule-specific PLAT domain genes in tissue responding to Rhizobia. A) Expression patterns of indicated genes at each timepoint by FPKM. Note the scale is calculated to a local maximum, indicated in red, which varies by the gene. B) Average expression, with error bars of the same genes in B in whole root segments under the same conditions (from data in Schnabel, et al 2023).

### Rhizobial response genes and their expression patterns

Because the LCM data set was generated from inoculated roots, we expected that in addition to genes involved in nodulation, genes involved in root development but not nodulation would also change expression over time. In order to focus on nodulation regulatory genes when examining temporal patterns, we used the data from (Schnabel et al., 2023) to identify genes showing an increase in expression in root segments responding to rhizobia in wild type and/or in the autoregulation of nodulation (AON) mutants *sunn-1* and *rdn1-2* when compared to uninoculated A17 controls; we termed this collection of genes “rhizobial response genes” (n=1932; Supplemental Data Set 2A). All but two of the genes identified as differentially expressed in wild type root segments responding to rhizobia were also identified as differentially expressed in AON mutants responding to rhizobia; thus the AON set appears to be an extension of the wild-type set (Supplemental Figure 2).

A large percentage of the rhizobial response genes (82.7%) were previously identified as differentially expressed in nodulating roots for at least one time point (2 to 72 hpi) in a study using similar conditions, ecotypes and rhizobia (Schiessl et al., 2019). The overlapping genes included 82.5% of the genes detected in the wild-type set and 82.7% of the genes detected in the AON set. Beyond simple differential expression at a single time point, we sorted the rhizobial response genes into groups by temporal pattern of expression and created a heat map to display the distribution of expression patterns (Figure 5). Genes were classified and placed in groups based upon time of the first increase in expression from the level at 0 hpi (Early, I; Middle, II; or Late, III in column 3 of Supplemental Data Set 2A). We then further classified the list based on the subsequent pattern of this increase over our time course. If the increase was transient and later decreased, it was assigned the label “down” (A in Figure 5). If expression increased and persisted at the same level the label is ‘stays” (B in Figure 5) and if the expression level of the gene continued to increase over the time course the label was “increases” (C in Figure 5). In this classification, genes that increased expression from 0 hpi only at 72 hpi are simply labeled “Late” as there is no pattern to assign. The resulting categories are Early Down (IA), Early Stays (IB), Early Increases (IC), Middle Down (IIA), Middle Increases (IIB), Middle Stays (IIC), and Late (III).

**Figure 5.**
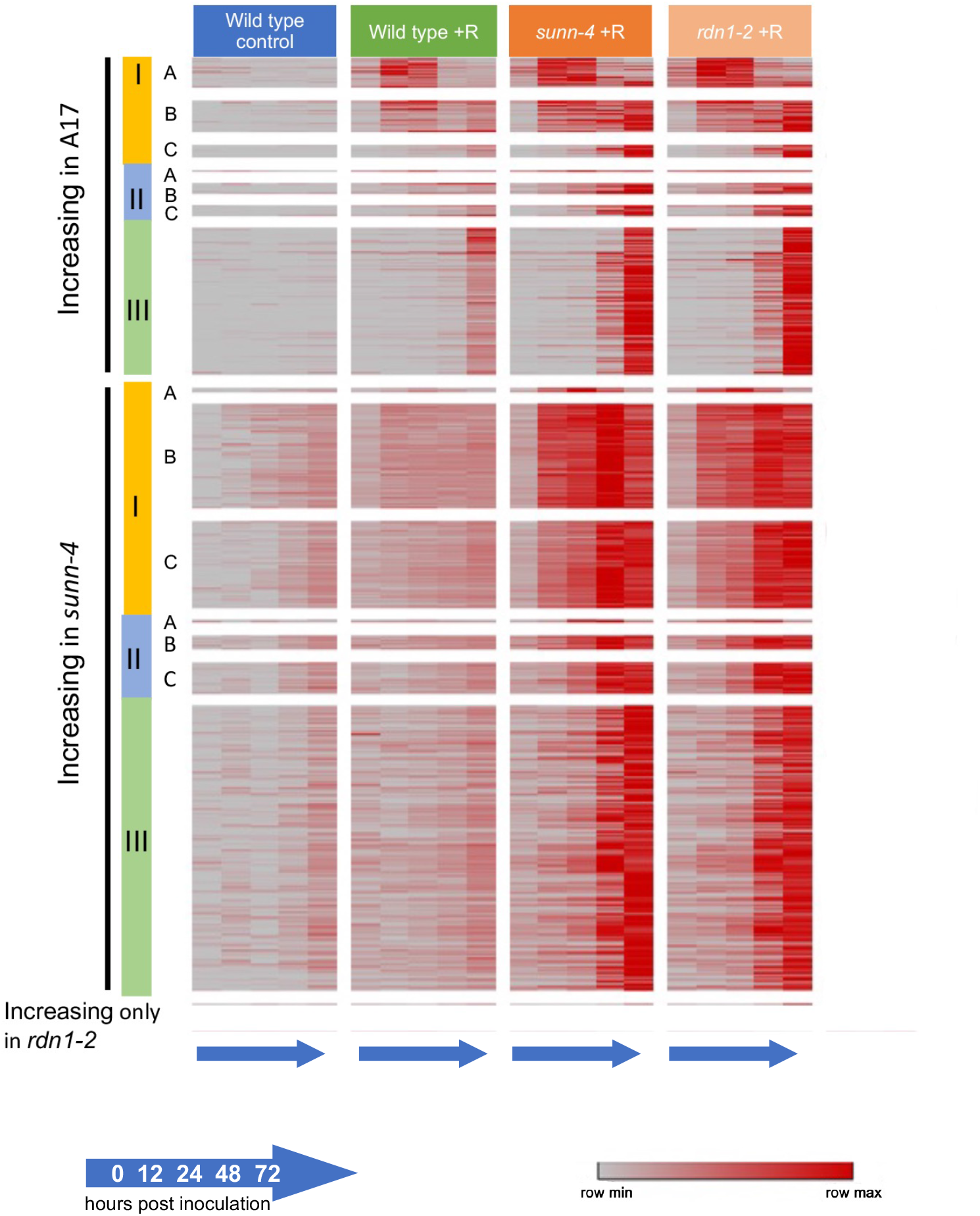
Rhizobial response genes show several different temporal expression patterns in root segments. Heat map of the 1932 rhizobial response genes identified from (Schnabel et al., 2023) and sorted by temporal pattern of expression. Shown are genes that increased in A17, (the majority of which also increased in *sunn-4* and *rdn1-2)*, increased in *sunn-4* but not A17 (including genes that also increased in *rdn1-2)*, or increased in *rdn1-2* only (one gene). Yellow (I), blue (II) and green (III) refer to Early, Middle, and Late time points of first observed increase in expression (see text for definition). Letters down the y axis refer to a pattern of expression defined as A, transient (up and then decreasing); B, persistent (up and holding steady); and C, increasing (steady increase throughout).

### Comparison of Root Segment expression patterns to the Laser Capture Data set

Armed with a gene set exhibiting patterns of temporal response to rhizobia in root segments, we performed a manual curation analysis of the LCM data with the categories defined above. The defined temporal expression patterns (DTEPs) of the rhizobial response gene set identified in segments of roots could be driven by the same DTEP for a gene being present in one or more tissues or it could be the result of summing the different DTEPs across all tissues. To focus on which expression patterns in each tissue were most important to early nodulation signaling, we compared DTEPs of the rhizobial response gene set generated from root segments in Figure 5 to the DTEPs seen in individual tissues in the LCM data set (see Materials & Methods for description of the curation). Initial analysis determined that data from nodule tissue at 48 and 72 hours drove the assignment of DTEPs and masked early signaling in other tissues. Therefore, we removed the nodule tissue and timepoint data, and analyzed the other tissue and timepoint data separate from the nodule tissue data while normalizing expression within each remaining tissue.

In the remaining tissues, 417 of the 1932 rhizobial response genes identified as having a DTEP in root segments displayed one of the DTEPs in at least one tissue. (Table 1; Supplemental Data Set 2B). The following analysis does not include the 43% of genes (842) from Supplemental Data Set 1A that changed in response to rhizobia but did not show one of the defined DTEPs in individual tissues versus in intact root segments. Taken together, 1090 DTEPs in a tissue were identified; many genes displayed the same pattern in multiple tissues, and some displayed different patterns in different tissues. Only 21% of these genes (102) had a single DTEP in a single tissue (marked in yellow in Supplemental Data Set 2B). Five genes began increasing at 12 or 24 hpi and continued to increase in a specific tissue (“Increases” category) compared to 46 genes in this category in the whole root segments (Category II in Figure 5). For simplification of the analysis, the “Stays” and “Increases” categories are combined in Table 1, however the 5 genes originally assigned to “Increases” are still marked as such in the Supplemental Data Set 2B to maintain the original data.

**Table 1.** Rhizobial Response genes identified as having a DTEP in root segments.

| Time of 1<sup>st</sup><br>increase | Early<br>Down | Early<br>Stays/Increases | Middle<br>Down | Middle<br>Stays/Increases | Late | Total<br>w/pattern | Tissue<br>specific |
| --- | --- | --- | --- | --- | --- | --- | --- |
| Tissue |  |  |  |  |  |  |  |
| ED | 77 | 45 | 67 | 49 | 82 | 320 | 53 |
| OC | 89 | 107 | 36 | 26 | 34 | 292 | 37 |
| ICA | 36 | 109 | 10 | 38 | 16 | 209 | 4 |
| ICB | 22 | 30 | 12 | 25 | 20 | 109 | 0 |
| VS | 33 | 42 | 24 | 22 | 39 | 160 | 9 |
| **Total:** | **257** | **333** | **149** | **160** | **191** | **1090** | **102** |

A broad overview of the refined gene set in Supplemental Data Set 2B examined by tissue showed that more than half the genes with a DTEP in a tissue showed that pattern in either the epidermis or the outer cortex (ED-28.5%; OC-26%) (Figure 6A). Cells in these tissues are the first to physically contact the rhizobial infection thread, which proceeds from the epidermis into the cortex. The remaining 44% had a DTEP in inner cells with this breakdown: the inner cortical cells at the xylem poles (ICA-18.6%), the inner cortical cells between the xylem poles (ICB-9.7%), and the vasculature cells (VS-14.2%). Of the 102 genes that had a DTEP in only one tissue, 87% were in the ED and OC cells (53 and 36 genes respectively-Figure 6B). There were also 33 genes that had a DTEP in every tissue, and these genes are highlighted in green in Supplemental Data Set 2B.

**Figure 6.**
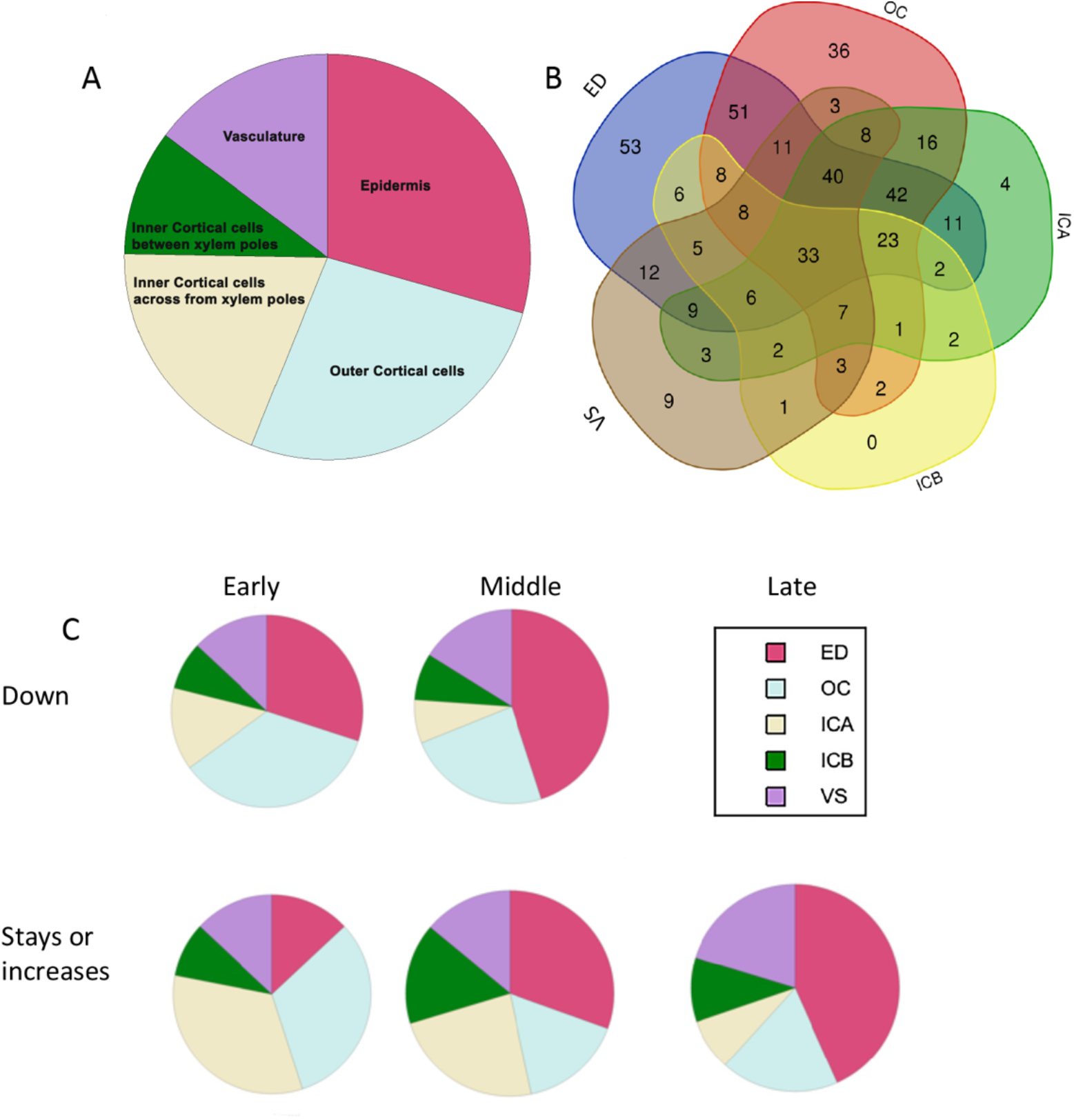
Distribution of rhizobial response genes with DTEPs between tissues. **See** text for description of the DTEP patterns (A) Pie chart indicating percentage of the rhizobial response Genes identified with a pattern in each tissue. (B) Venn Diagram of sharing of gene patterns between tissues. The number shared genes between tissues are indicated in each overlap, with tissues indicated by color. ED=epidermis (blue), OC=outer cortical cells (red), ICA=inner cortical cells across from xylem poles (green), ICB=inner cortical cells between xylem poles (yellow) and VS=vasculature (tan). (C) Pie charts showing distribution of patterns by tissue. Raw data is in Table 1 and Table 2.

Comparing the DTEPs to each other without including the nodule tissue (Table 2), over half the patterned genes (54%) increased in at least one tissue by 12 hours after inoculation and another 28% by 24 hours (Figure 6C). Only 17% increased at 48 hours or later. Note that this analysis did not include the genes that increased only in nodule tissue, which by our definition would always be considered as “Late.” Interestingly, 43% of the late genes in our subset were in the epidermal tissue suggesting the epidermis is still transcriptionally responding to rhizobia even though infection threads have formed and cells have begun dividing to make nodules.

**Table 2.** Distribution of Rhizobial Response Genes with DTEPs by Time and Tissue

| Pattern:<br>Tissue (below) | Early<br>Down | Early<br>Stays/Increases | Middle Down | Middle<br>Stays/Increases | Late |
| --- | --- | --- | --- | --- | --- |
| ED | 30% | 13% | 45% | 31% | 43% |
| OC | 35% | 32% | 24% | 16% | 18% |
| ICA | 14% | 33% | 7% | 24% | 8% |
| ICB | 8% | 9% | 8% | 16% | 10% |
| VS | 13% | 13% | 16% | 14% | 20% |
| Total # of genes | 257 | 333 | 149 | 160 | 191 |
| **Percent of total** | **EARLY** | **54%** | **MIDDLE** | **28%** | **LATE 17%** |

Rhizobial response genes that were expressed in nodules at 48 and 72 hpi were examined separately. Many of these genes were also expressed in other tissues, but a subset was expressed only in nodules. Sixty percent (66 genes) of the 110 genes in the “Late” increase category of our initial temporal pattern analysis were exclusively expressed in nodule tissue and are listed in Supplemental Data Set 2C.

### Genes that influence the spatial distribution of nodules

As an example of using our LCM data set to address questions about early nodulation signaling, we attempted to identify the signaling events that result in some inner cortical cells attracting infection threads to become nodules while other cells remain in the G0 cell cycle stage. While the observation of a local difference in ethylene concentration based on the differential expression of a 1-aminocyclopropane-1-carboxylate oxidase (ACC oxidase) gene between legume inner cortical cells in relation to their position relative to the xylem poles was made over 25 years ago (Heidstra et al., 1997), the transcriptional changes that make these cells different remain unknown. Specifically, in *Pisum sativum*, mRNA encoding one of many ACC oxidases was shown by staining to accumulate in the cell layers opposite the phloem poles (ICA) at 6 days post inoculation (Heidstra et al., 1997). The *M. truncatula sickle* mutant, which lacks regulation of nodule position, has lesions in the ortholog of *EIN2* involved in ethylene signaling (Penmetsa and Cook, 1997; Penmetsa et al., 2008). Together these data implicate ethylene in spatial positioning.

We therefore visualized the expression of all *M. truncatula* ACC oxidase genes across our cell types during early nodule development in the LCM data set, including the putative ortholog of the pea ACC oxidase gene (Medtr2g025120) used in the 1997 study (Peck et al., 1993; Heidstra et al., 1997) (Figure 7). This transcript and two others (Medtr1g043760 and Medtr6g092620) had unequal expression between the ICA and ICB at zero hours in the absence of rhizobia (Figure 7). At 12 hours, all ACC oxidase genes except Medtr2g025120 and Medtr5g085330 were expressed at the same relative level in both the ICA and ICB. Both Medtr2g025120 and Medtr6g092620 demonstrated clear expression differences in ICA versus ICB at 12 and 24 hpi respectively while Medtr5g085330 showed a difference between ICA and ICB cells only after the nodules formed at 72 hpi. Thus, all four of the ACC oxidases were expressed more highly in the ICB than the ICA at some point in the nodulation time course, suggesting this expression difference is a possible marker for cells that will not become nodules. Because we saw nodule cell division in our system at 48 hours (Figure 1A), a comparison of gene expression in the tissue where most nodules will form (ICA) against where they mostly are excluded (ICB) at earlier time points could identify transcriptional differences beyond ethylene production at the early timepoints.

**Figure 7:**
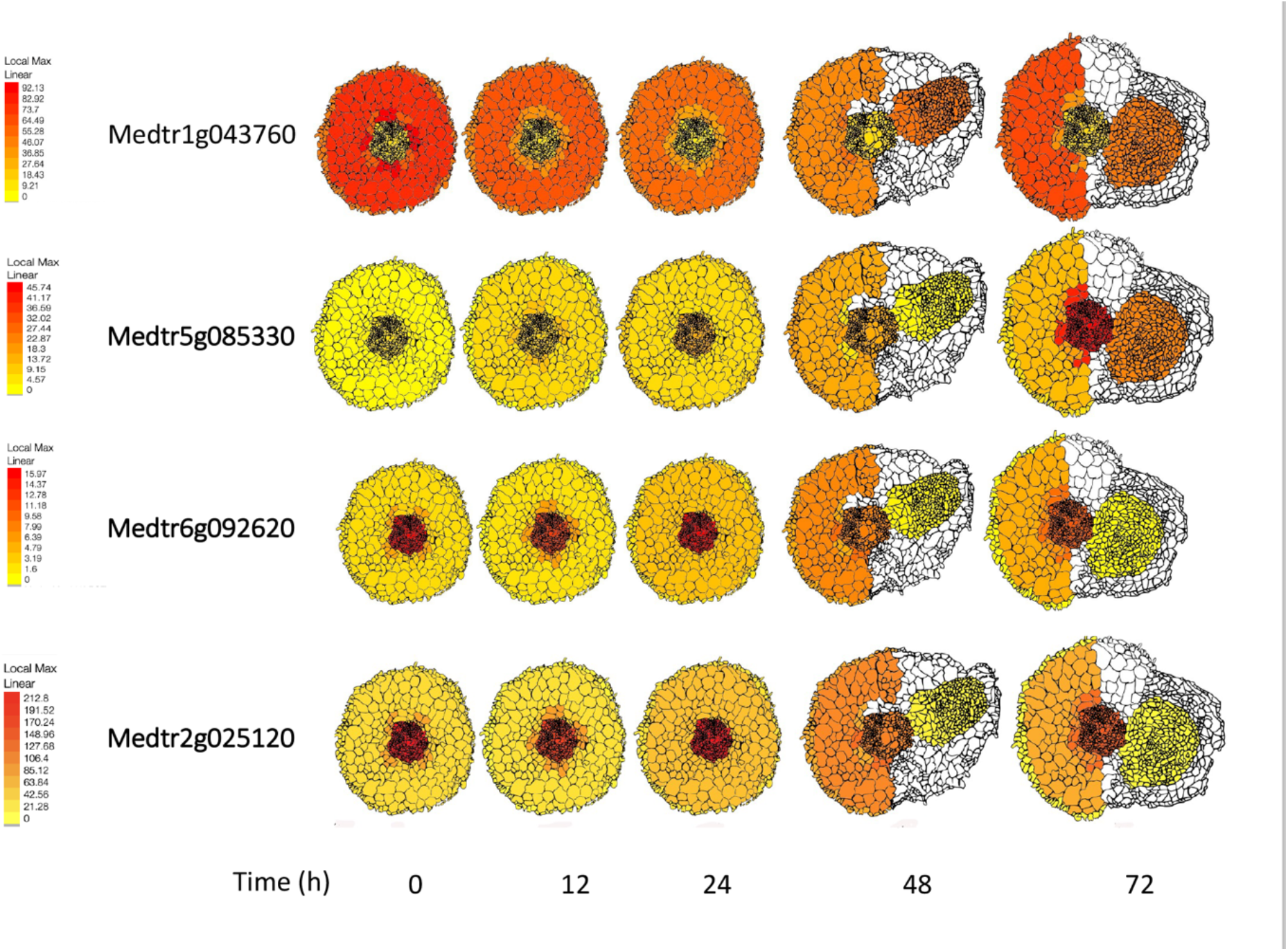
Tissue Expression of *M. truncatula* ACC synthases in early nodulation signaling. Expression patterns of indicated ACC synthase genes at each timepoint by FPKM. Note the scale is calculated to a local maximum, indicated in red, which varies by the gene.

Thus, we applied two non-overlapping approaches (DeSeq2, GeneShift) to identifying potential genes specific to the spatial position of the inner cortical cells that become nodules. The first approach used DeSeq2 (Love, et al. 2014) to identify differential gene expression at single time points and identified all genes in the ICB and ICA tissues indicated by DeSeq2 as a differentially expressed between 0 and 12 hpi or 0 and 24 hpi. The list was curated to remove genes with very low FPKMs, for which the overall pattern in the inner cortical cells from 0 to 72 was probably noise, under the assumption the differential expression at one timepoint in a single replicate would be an anomaly and any gene with a similar pattern between the two tissues even though only one tissue was flagged by DeSeq2. Heat maps of genes induced early in the ICB but not the ICA (32 genes) and genes induced early in the ICA but not the ICB (30 genes) are presented in Supplemental Figure 3; the lists and gene annotations are found in Supplemental Data Set 2D.

We compared these lists to the rhizobial response genes list generated above and discovered three rhizobial response genes induced early in the ICB but not the ICA (Medtr5g056160, an S1/P1 nuclease family protein; Medtr2g007990, an auxin-induced 5NG4-like protein; and Medtr1g011650, an ABC-2 and plant PDR ABC-type transporter family protein). We also identified four rhizobial response genes induced early in the ICA but not the ICB (Medtr7g074730, a dihydroflavonol 4-reductase-like protein; Medtr4g067090, a hypothetical protein; Medtr6g014270, a UDP-glucosyltransferase family protein; and Medtr4g055580, an F-box/RNI/FBD-like domain protein) all highlighted in Supplemental Data Set 2D. Testing of the entire set for functional enrichment revealed statistical enrichment (p<0.05) for hydrolase activity, transporter activity, plasma membrane and “other” intracellular components in genes that were induced early in the ICB but not the ICA, and only for plastid, and “other” cellular and intracellular components, in genes induced early in the ICA but not the ICB (Figure 8A). Applying a test for enrichment of gene families (Bedre and Mandadi, 2019) to the list of genes induced only in cells where the nodules form (ICA) revealed that gene families for nodulin-like genes, glucan synthase-like genes, alpha-galactosidase genes, the PPIase and Dof gene families are overrepresented (Figure 8B). In cells where the nodules do not form (ICB), induced gene families for pentatricopeptide repeat genes, glutaredoxins, caffeic acid o-methyltransferases, and thioredoxins are overrepresented (Figure 8B).

**Figure 8:**
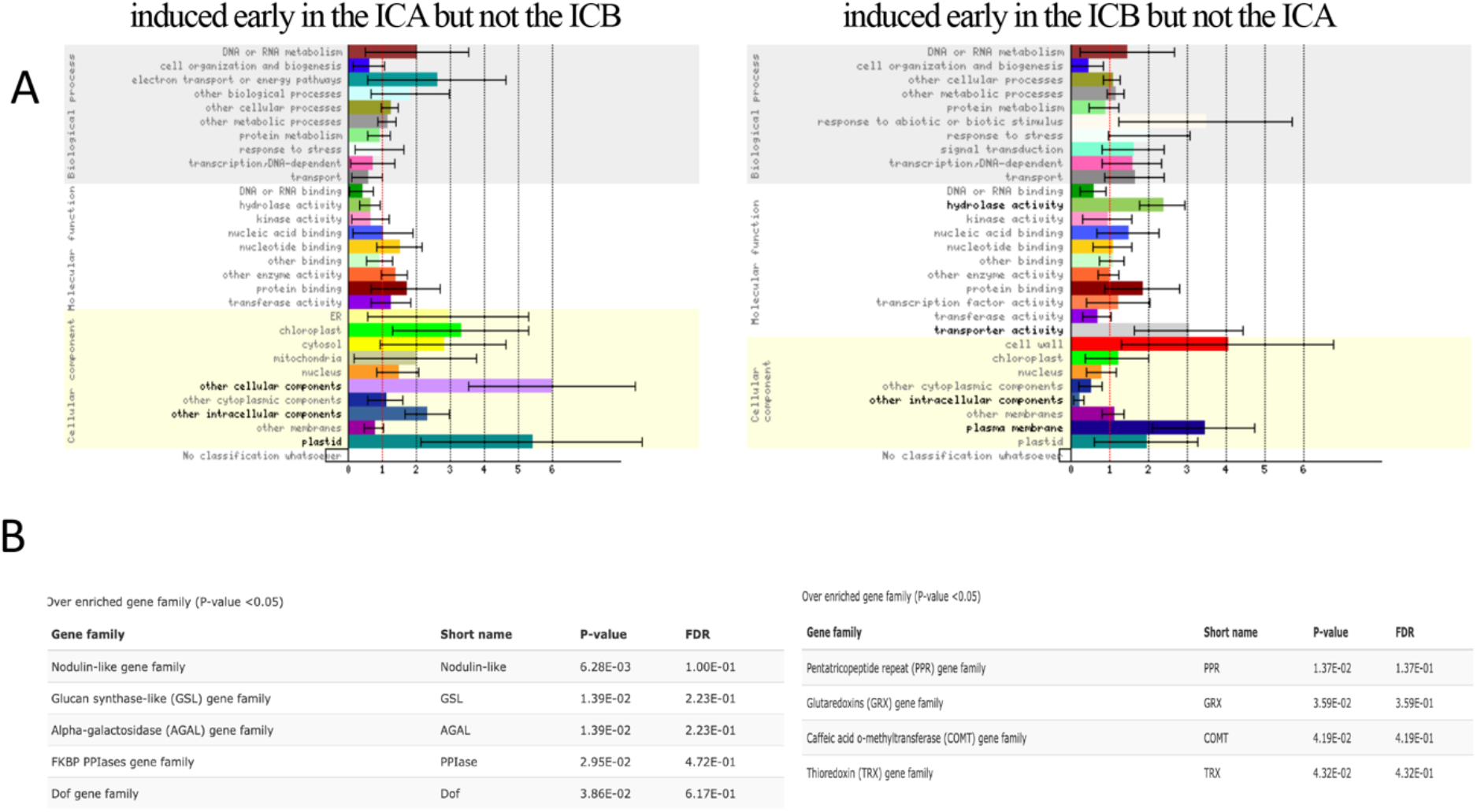
Enrichment of lists of genes differentially expressed between the ICA and the ICB. (A) GO term functional enrichment of lists from Supplemental Dataset 2D using Classification Super Viewer tool (see Methods). GO categories are displayed for each GO subclass ranked by normed frequency values. Errors bars indicate the standard deviation of the normed frequency. Terms with statistical enrichment determined by Hypergeometric enrichment tests (p<0.05) are marked with an asterisk. (B) Gene families over-enriched in each category according to GenFam (p <0.05), which uses a Fisher Exact test with a Bonferroni corrected p-value.

In a second approach we used GeneShift (Gao et al., 2022) to analyze differing patterns of genes expressed in the ICA and ICB over the entire time course without the step of defining DTEPs of genes and focused on any pattern differences during 0-24 hpi. GeneShift identifies genes with statistically similar patterns of expression, especially those that are expressed at low levels; these genes were eliminated in the first analysis because of the parameters of DeSeq2. GeneShift outputs trajectory sets, grouping genes with a similar pattern defined by the algorithm as statistically significant (Gao et al., 2022). GeneShift identified 52 genes in 9 trajectory sets with different patterns of expression in the two tissues after inoculation (Supplemental Data Set 2E). Genes in category A (23 genes in 5 trajectory sets) demonstrated dynamic expression in ICA, while in ICB at least one biological replicate recorded expression levels of 0.0 across all harvested time points. Genes in category B (29 genes in 4 trajectory sets) demonstrated no differential expression in at least one biological replicate in ICA but showed dynamic expression in ICB. The heat map of the individual replicates is shown in Supplemental Figure 4. These trajectory sets did not use the differential expression criteria used in creating the rhizobial response gene set but did compare pattern differences between the inner cortical cell tissues responding to rhizobia, resulting no overlap between the set of genes from the first analysis. While samples of the patterns identified for all trajectory sets are displayed in Supplemental Figure 5, we chose to focus on the trajectory sets with pattern differences between ICA and ICB cells at the 0, 12 (A5, B4), and 24 (A4, B3) hpi before nodule cell division begins.

In the ICA tissue, the A2 trajectory set contains nine genes with the same pattern of expression differences between ICA and ICB tissues at 0 hpi, suggesting a functional difference between the two cell types prior to rhizobia inoculation (Supplemental Figure 5). One of these genes, Medtr6g009090, an ABC transporter transmembrane protein, is also on the rhizobial response gene list, while the others (a transmembrane protein, as well as disease resistance genes, transcription factors, a receptor kinase, and others) are not. The A5 trajectory set containing genes with dynamic expression in the ICA tissue at 12 hpi had only two genes: Medtr1g029070, which encodes the NF-YB transcription factor and Medtr6g074780, which encodes a disease-resistance gene. The A4 trajectory set genes with dynamic expression in the ICA at 24 hpi contains four genes, including the disease resistance gene *Rpp4C1* (Medtr8g028500) and a plant resistance-related chalcone synthase gene (Medtr5g007740).

In ICB cells where nodules do not form, trajectory sets B3 and B4 show dynamic expression in ICB tissue but not ICA tissue. B4 contains four genes with dynamic expression at 12 hpi, two of which are hypothetical proteins. B3 contains six genes with dynamic expression at 24 hpi, including two hypothetical proteins and a PPR-containing plant like protein (Medtr4g023510). Together the two approaches created gene lists that can now be tested for functional relevance and for transcriptional response to ethylene.

## Discussion

### Individual gene expression patterns

Our results show that a similar temporal pattern of the early nodulation genes in whole root segments was revealed by LCM as differential gene patterns in individual tissues (epidermis, outer cortex, inner cortex between the xylem poles, inner cortex across from xylem poles, and the vasculature). *VPY, NSP1*, and *NFH1* all show peak expression in the epidermal cells at 12 hours, but only *VPY* and *NSP1* also have a peak in nodule tissue. *NFH1* encodes the Nod factor hydrolase required for infection (Tian et al., 2013) and is induced in root hairs of soybean and *M. truncatula* in this time frame (Libault et al., 2010; Breakspear et al., 2014), thus confirming our results. *VPY* is part of a complex required for the polar growth of infection threads (Liu et al., 2019a). Increased expression of *VPY* in the epidermis during the first 24 hpi, followed by an upregulation in the developing nodule reflects its function in the polar growth of infection threads, as these are the entry (epidermis) and end points (nodule) of infection thread growth. *NSP1* is required for downstream signaling after Nod factor perception; *NSP1* with *NSP2* binds to *cis* regulatory elements of *ENOD 11, ERN1* and *NIN* genes (Oldroyd and Long, 2003; Kalo et al., 2005; Vernie et al., 2015; Liu et al., 2019b;(Roy et al., 2020; Liu et al., 2022). *NIN* is a central regulator of nodulation and known to play different roles in the epidermis versus the cortex. In addition, *NIN* and *ERN1* compete for *cis* regulatory elements on *ENOD11* (Vernie et al., 2015). This was also observed in the time course expression patterns in Figure 2B where *NIN* and *ERN1* have similar expression patterns (not intensity levels) in the epidermis at 0 hpi and in the cortex at 12 hpi. At 24 and 48 hpi, the expression patterns for *NIN* and *ERN1* were similar except that at 48 hpi, *ERN1* rose in the epidermis while *NIN* did not. At 72 hpi *NIN* and *ERN1* were both highly expressed in the nodule. Without the resolution provided by the spatio-temporal analysis showing only *ERN1* rising the epidermis at 48 hpi, the genes would have incorrectly appeared to be co-expressed.

Spatial patterns of peptide gene expression are more complex to interpret, since the CLE peptides are significantly post translationally processed (reviewed in Shinohara and Matsubayashi, 2010) and move through the plant to regulate symbiosis (reviewed in Roy and Müller, 2022). Nevertheless, localized expression patterns showing where the genes are induced, versus where the protein products act, can be useful for constructing regulatory pathways. For example, *MtCLE12* and *MtCLE13* are both required in AON, but can be distinguished by the requirement of the *RDN1* enzyme to process *MtCLE12* for functionality (Kassaw et al., 2017). Both genes have similar whole root expression patterns at 0 and 72 hpi, differing in the timing of induction between these points, but not the level of expression (Figure 3A). However, our tissue level analysis (Figure 3B) revealed that *MtCLE13* expression increased in cortical cells at 12 hpi followed by the vasculature and then the nodule, while *MtCLE12* expression did not increase until 48 hpi, only in the vasculature, followed by the nodule, suggesting the two peptides have different roles and/or targets. A similar conclusion can be drawn for the nodule-specific PLAT domain genes in Figure 4, which all exhibited strong expression in nodules. Since *MtNPD2* expression showed a secondary expression maximum in outer cortical cells at 24 hpi, and *MtNDP3,4* and *5* all showed a secondary maximum in inner cortical cells at 24 hpi, the genes might be better termed nodule-associated as opposed to nodule-specific. Only *MtNDP1* appears to be truly nodule specific and *MtNDP2* could have a different function than *MtNDP3,4* and *5* given they have different spatiotemporal expression patterns.

### Global tissue response to nodulation signals

By looking at patterns of temporal response instead of simply determining which genes increase expression during or after nodulation, some global observations are now possible regarding which tissues are responding to rhizobia. First, increased gene expression in the developing nodules at 48 and 72 hpi dominated our analysis. Second, 77.5% of the genes with expression in nodules were also expressed in other tissues -- there were only 66 “nodule specific” genes at this point in development which is in line with the theory that nodules are modifications of the lateral root development plan (Schiessl et al., 2019; Soyano et al., 2021; Schiessl et al., 2023). Therefore, nodule specific genes are not likely to be involved in early nodulation signaling (12-24 hpi). Third, when nodule tissue was removed from analysis, not surprisingly over half the genes with a DTEP were in the epidermis and the cortical cells, which undergo changes that allow the rhizobia to enter the plant through the infection thread, and the infection thread to pass through to the inner cortical cells that will become the nodule. Our DTEP gene lists provide starting points for investigations of nodulation signaling between tissues. These gene lists can also be used to confirm cell to gross level tissue assignments in single-cell RNASeq experiments, since the tissues are determined by physical capture versus informatic analysis.

### Identifying genes involved in nodule positioning

The expression patterns of ACC oxidases provide further evidence that ethylene is a signal for the spatial positioning of nodules, but the identification of potential downstream transcriptional responses to the ethylene signal also provides a starting point for further investigation into the nodulation process. The genes and gene families we identified in the tissue where nodules initiate (ICA) include disease resistance and nodulin-like genes as well as gene families involved in sugar synthesis. This suggests that by 24 hpi the cells destined to become nodules are already responding transcriptionally and is evidence that the signal for cell division is very early and may be perceived only in cells with low ACC synthase transcription, a differential gene expression observed at 0 hpi, which is before rhizobial inoculation. This observation is consistent with the phenotype of the ethylene insensitive *skl* mutant-nodules forming randomly in the inner cortical cells (Penmetsa and Cook, 1997; Penmetsa et al., 2003; Penmetsa et al., 2008).

In summary, the LCM time course data set we created allows examination of the spatiotemporal expression pattern of any of 4,893 *M. truncatula* genes during the early stages of nodule development. We characterized nodulation gene lists by examining gene expression between tissues and between uninoculated and rhizobial treatments. We also detected expression differences between tissue and conditions at single time points and across the full time series. Finally, the ePlant visualization and data-mining resource allows for public queries of our data to facilitate further discovery of the molecular control of the nodulation process.

## Materials and Methods

### Tissue capture

Tissue at five timepoints in three biological replicates of 10 plants each was obtained as described in (Schnabel et al., 2023) and fixed immediately as described in (Chavan et al., 2018). Fixed blocks of root segments were stored at -80°C until dewaxed and sectioned as described in (Chavan et al., 2018). For consistency, individual tissues to generate the libraries used were captured by only three authors (R.E., J.T. and S.C.) using the same internal lab protocol and equipment. After the experiments were finished and data was compiled and analyzed, one library (epidermis at 48 hpi) was incorrectly labeled based on comparison of the gene expression profile with other tissues at 48 hpi and was removed from the analysis, leaving 74 libraries.

### RNA prep, libraries, and sequencing

RNA was purified from captured sections using the Ambion RNAqueous-Micro Total RNA Isolation Kit (Invitrogen). Quality and concentration of RNAs were assayed on a 2100 Bioanalyzer (Agilent) using the RNA 6000 Pico kit. cDNAs were prepared from 1 ng of total RNA using the SMART-Seq v4 Ultra Low Input RNA Kit for Sequencing (TaKaRa) according to the manufacturer guidelines for cDNA synthesis and amplification, using 11 to 13 cycles of PCR amplification. The cDNAs were purified using AMPure XP beads as indicated in the SMART-Seq kit, eluting with the recommended 17 ml of elution buffer. cDNA quality and quantity were assayed on a 2100 Bioanalyzer using the High Sensitivity DNA kit. Libraries were prepared from 250 pg of cDNA using the Nextera XT DNA Library Preparation Kit (Illumina) according to the manufacturer’s guidelines using i7 and i5 indices. Libraries were assessed using the Agilent High Sensitivity DNA kit. Sequencing was performed by Novogene, with read pair counts per library ranging from 32,568,135 to 320,892,877 with a median of 71,070,324. Reads were mapped to v4.2 and v5.0 of the *M. truncatula* genome for the ePlant resource; ver 4.2 assignments are used in this manuscript.

### Data Availability

Data from LCM libraries was deposited in BioProject PRJNA704996. Data from root segment libraries is from BioProject PRJNA524899 (control A17) and BioProject PRJNA554677 (nodulating A17, *sunn-4*, and *rdn1-2* and control *sunn-4*). The data are also available in the Medicago ePlant at https://bar.utoronto.ca/eplant_medicago/.

### Manual curation of rhizobial response patterns (DTEPs)

The 1932 rhizobial response gene list from Figure 5 was curated for patterns in individual tissues by first removing any genes that met either of two noise criteria: 1) any gene with an assigned pattern driven by only one of the three replicates was removed, 2) any gene with all replicate FPKM values for a given tissue ≤l FPKM in that tissue was eliminated from consideration. The individual tissue expression pattern of the remaining genes was then normalized with a log2 transformation of all FPKM values, followed by hierarchical clustering of each gene by pattern, with each tissue treated separately and visualized by a Morpheus heatmap (https://software.broadinstitute.org/morpheus). Any gene that did not visually display one of the previously identified expression patterns (IA, IB, IC, IIA, IIB, IIC, III) in at least one tissue was removed from the analysis (289 genes).

For the 706 genes remaining, within each tissue replicates were averaged, and fold change calculated relative to the previous time-point (i.e., 12-hpi (A) calculated relative to 0-hpi time (B), the 24-hpi to the 12-hpi, etc.). The calculated fold changes were then used to determine if the expression of a gene increased (a ≤ 2.0 FC between A and B) or decreased (a ≤0.5-fold change between A and time-point B). Patterns meeting the fold change cutoff were considered strong enough to confidently assign a pattern to the gene in that tissue, and those not meeting the cutoff were removed. The final data set contained 417 genes.

Expression patterns in individual tissues were compared using a Venn Diagram analysis (http://bioinformatics.psb.ugent.be). For this by-tissue comparison, the genes were classified as responding Early (IA, IB, IC), Middle (IIA, IIB, IIC), or Late (III), as defined in the results, with Nodule tissue not considered, because nodule patterns are Late by definition.

### Inner Cortical cell analysis (DeSeq2 and GeneShift)

ICA and ICB data sets (Supplemental Figure 3) were processed from the raw data in Supplemental Dataset 1 with a DESeq2 pipeline using *Medicago truncatula* genome v4.2. The prepDE.py script from the StringTie Package [https://ccb.jhu.edu/software/stringtie/dl/prepDE.py] was utilized to compute the raw gene counts.

The DESeq2 R package (Love, et al. 2014) was employed to conduct the differential expression analysis, which has an internal normalization approach for the library size. Genes with a total read count of fewer than 50 were excluded from the analysis. The DESeqDataSetFromMatrix function was used to compare the 0h and 12h (or 24h) inoculated samples in ICA and ICB tissues, with the following formula: design = ∼ condition. Genes with an adjusted p-value of less than 0.05 were deemed significant. The same data was also processed with GeneShift (Supplemental Figures 4 and 5), a computational workflow for time-series gene expression profiling using the two tissues as conditions. GeneShift uses a six-stage process incorporating a deep learning classification model to obtain high quality expression pattern shifts between multiple conditions and timepoints and is described in (Gao et al., 2022).

### Enrichment Analysis

The graphical summary of the gene ontology (GO) classification (Figure 8) was performed with the Classification Super Viewer tool from http://bar.utoronto.ca_adapted to *M. truncatula* using the default settings (Herrbach et al., 2017). The data are the normed frequency of each GO category for the given sets of genes compared to the overall frequency calculated for the Mt4.0 *M. truncatula* genome. The lists of overrepresented gene families were created with GenFam, (https://www.mandadilab.com) (Bedre and Mandadi, 2019).

### Creating two new views for the Medicago ePlant

Scalable Vector Graphic (SVG) images describing the LCM tissue we captured as well as the root bulk tissue sections as described in (Schnabel et al., 2023) were created in InkScape (inkscape.org) based on micrographs similar to those shown in Figure 1 and sketches of longitudinal views of the sampled roots. XML files were manually created that associated labeled group tags in the SVG files with databased sample names plus expression data (see original ePlant publication Waese et al., 2017) and added to the Medicago ePlant instance (Waese-Perlman et al., 2021) running on the Bio-Analytic Resource for Plant Biology at http://bar.utoronto.ca as a Root Component view and a Root view, respectively.

## Supporting information

Supplemental Data Set 1

Supplemental Data Set 2

## Supplemental Figures

Supplemental Figure 1. Diagram of procedure designed to increase signal to noise ratio

Supplemental Figure 2. Overlap of rhizobial response genes between wild type and supernodulation mutants.

Supplemental Figure 3. Genes with differential expression in spatially differentiated inner cortical cells at early timepoints

Supplemental Figure 4: Gene Expression Profiles in ICA and ICB in inoculated *M. truncatula* over 72 Hours.

Supplemental Figure 5. Expression profiles of representative genes in GeneShift trajectory sets for ICA and ICB tissues.

## Supplemental Data Files

Supplemental Data Set 1. FPKM values of all genes detected in all libraries

Supplemental Data Set 2:

2A. Rhizobia response genes in nodulating root segments of *M. truncatula*

2B. Rhizobial response genes displaying temporal patterns in captured tissues of *M. truncatula*

2C. Rhizobial response genes displaying temporal patterns in nodule tissue of *M. truncatula*

2D. *M. truncatula* genes identified by DeSeq2 as having different patterns of expression in inner cortical cells across from the xylem poles (ICA) and between the xylem poles (ICB) at early time points (0-24 hours post inoculation)

2E. *M. truncatula* genes identified by Gene Shift as having different patterns of expression in inner cortical cells across from the xylem poles (ICA) and between the xylem poles (ICB) at early time points (0-24 hours post inoculation)

## Author Contributions and Acknowledgements

ES, JT, RE, YG, SC, WP, and JF performed experiments, mapped data sets, and created or contributed to creation of figures. AP, EE, and NP created the ePlant interface, JF and FAF obtained the funding. All authors wrote and edited the manuscript. This work was supported by NSF IOS 1444461 to JF and FAF and NSF IOS 1733470 to JF. Images and tissue were captured using the Clemson Light Imaging Facility and the work was made possible, in part, with support from the Clemson University Genomics and Bioinformatics Facility which receives support from an Institutional Development Award (IDeA) from the National Institute of General Medical Sciences of the National Institutes of Health under grant number P20GM109094. Much of the computation was performed on the Clemson University Palmetto Cluster.

## Literature Cited

Bedre, R., and Mandadi, K. (2019). GenFam: A web application and database for gene family-based classification and functional enrichment analysis. Plant Direct 3, e00191.

Bhuvaneswari, T., Turgeon, B.G., and Bauer, W.D. (1980). Early events in the infection of soybean (Glycine max L. Merr) by Rhizobium japonicum: I. Localization of infectible root cells. Plant Physiology 66, 1027–1031.

Breakspear, A., Liu, C.W., Roy, S., Stacey, N., Rogers, C., Trick, M., Morieri, G., Mysore, K.S., Wen, J.Q., Oldroyd, G.E.D., Downie, J.A., and Murray, J.D. (2014). The Root Hair “Infectome” of Medicago truncatula Uncovers Changes in Cell Cycle Genes and Reveals a Requirement for Auxin Signaling in Rhizobial Infection. Plant Cell 26, 4680–4701.

Brewin, N.J. (1991). Development of the legume root nodule. Annu Rev Cell Biol 7, 191–226.

Cai, J., Veerappan, V., Arildsen, K., Sullivan, C., Piechowicz, M., Frugoli, J., Dickstein, R. (2023) A Modified Aeroponic System for Growing Plants to Study Root Systems. Plant Methods, In press.

Capoen, W., Sun, J., Wysham, D., Otegui, M.S., Venkateshwaran, M., Hirsch, S., Miwa, H., Downie, J.A., Morris, R.J., Ané, J.-M., and Oldroyd, G.E.D. (2011). Nuclear membranes control symbiotic calcium signaling of legumes. Proceedings of the National Academy of Sciences of the United States of America 108, 14348–14353.

Cervantes-Pérez, S.A., Thibivilliers, S., Laffont, C., Farmer, A.D., Frugier, F., and Libault, M. (2022). Cell-specific pathways recruited for symbiotic nodulation in the Medicago truncatula legume. Molecular Plant 15, 1868–1888.

Charon, C., Johansson, C., Kondorosi, E., Kondorosi, A., and Crespi, M. (1997). enod40 induces dedifferentiation and division of root cortical cells in legumes. Proc Natl Acad Sci U S A 94, 8901–8906.

Chavan, S., Schnabel, E., Saski, C., and Frugoli, J. (2018). Fixation and Laser Capture Microdissection of Plant Tissue for RNA Extraction and RNASeq Library Preparation. Current Protocols in Plant Biology 3, 14–32.

Ehrhardt, D.W., Wais, R., and Long, S.R. (1996). Calcium spiking in plant root hairs responding to Rhizobium nodulation signals. Cell 85, 673–681.

Gage, D.J. (2004). Infection and Invasion of Roots by Symbiotic, Nitrogen-Fixing Rhizobia during Nodulation of Temperate Legumes. Microbiology and Molecular Biology Reviews 68, 280–300.

Gamas, P., Brault, M., Jardinaud, M.-F., and Frugier, F. (2017). Cytokinins in symbiotic nodulation: when, where, what for? Trends in Plant Science 22, 792–802.

Gao, Y., Selee, B., Schnabel, E.L., Poehlman, W.L., Chavan, S.A., Frugoli, J.A., and Feltus, F.A. (2022). Time Series Transcriptome Analysis in Medicago truncatula Shoot and Root Tissue During Early Nodulation. Frontiers in Plant Science 13.

Gaude, N., Bortfeld, S., Duensing, N., Lohse, M., and Krajinski, F. (2012). Arbuscule-containing and non-colonized cortical cells of mycorrhizal roots undergo extensive and specific reprogramming during arbuscular mycorrhizal development. The Plant Journal 69, 510–528.

Gomez, S.K., Javot, H., Deewatthanawong, P., Torres-Jerez, I., Tang, Y., Blancaflor, E.B., Udvardi, M.K., and Harrison, M.J. (2009). Medicago truncatula and Glomus intraradices gene expression in cortical cells harboring arbuscules in the arbuscular mycorrhizal symbiosis. BMC plant biology 9, 1–19.

Haney, C.H., and Long, S.R. (2010). Plant flotillins are required for infection by nitrogen-fixing bacteria. Proceedings of the National Academy of Sciences of the United States of America 107, 478–483.

Heidstra, R., Yang, W.C., Yalcin, Y., Peck, S., Emons, A.M., vanKammen, A., and Bisseling, T. (1997). Ethylene provides positional information on cortical cell division but is not involved in Nod factor-induced root hair tip growth in Rhizobium-legume interaction. Development 124, 1781–1787.

Herrbach, V., Chirinos, X., Rengel, D., Agbevenou, K., Vincent, R., Pateyron, S., Huguet, S., Balzergue, S., Pasha, A., and Provart, N. (2017). Nod factors potentiate auxin signaling for transcriptional regulation and lateral root formation in Medicago truncatula. Journal of experimental botany 68, 569–583.

Horvath, B., Yeun, L.H., Domonkos, A., Halasz, G., Gobbato, E., Ayaydin, F., Miro, K., Hirsch, S., Sun, J.H., Tadege, M., Ratet, P., Mysore, K.S., Ane, J.M., Oldroyd, G.E.D., and Kalo, P. (2011). Medicago truncatula IPD3 Is a Member of the Common Symbiotic Signaling Pathway Required for Rhizobial and Mycorrhizal Symbioses. Molecular Plant-Microbe Interactions 24, 1345–1358.

Huo, X., Schnabel, E., Hughes, K., and Frugoli, J. (2006). RNAi Phenotypes and the Localization of a Protein::GUS Fusion Imply a Role for Medicago truncatula PIN Genes in Nodulation. J Plant Growth Regul 25, 156–165.

Jardinaud, M.F., Boivin, S., Rodde, N., Catrice, O., Kisiala, A., Lepage, A., Moreau, S., Roux, B., Cottret, L., Sallet, E., Brault, M., Emery, R.J., Gouzy, J., Frugier, F., and Gamas, P. (2016). A Laser Dissection-RNAseq Analysis Highlights the Activation of Cytokinin Pathways by Nod Factors in the Medicago truncatula Root Epidermis. Plant Physiol 171, 2256–2276.

Jin, Y., Liu, H., Luo, D., Yu, N., Dong, W., Wang, C., Zhang, X., Dai, H., Yang, J., and Wang, E. (2016). DELLA proteins are common components of symbiotic rhizobial and mycorrhizal signalling pathways. Nature Communications 7, 12433.

Kalo, P., Gleason, C., Edwards, A., Marsh, J., Mitra, R.M., Hirsch, S., Jakab, J., Sims, S., Long, S.R., Rogers, J., Kiss, G.B., Downie, J.A., and Oldroyd, G.E. (2005). Nodulation signaling in legumes requires NSP2, a member of the GRAS family of transcriptional regulators. Science 308, 1786–1789.

Kassaw, T., Nowak, S., Schnabel, E., and Frugoli, J. (2017). ROOT DETERMINED NODULATION1 Is Required for M. truncatula CLE12, But Not CLE13, Peptide Signaling through the SUNN Receptor Kinase. Plant Physiol 174, 2445–2456.

Kassaw, T.K., and Frugoli, J.A. (2012). Simple and efficient methods to generate split roots and grafted plants useful for long-distance signaling studies in Medicago truncatula and other small plants. Plant Methods 8, 38.

Lefebvre, B., Timmers, T., Mbengue, M., Moreau, S., Hervé, C., Tóth, K., Bittencourt-Silvestre, J., Klaus, D., Deslandes, L., Godiard, L., Murray, J.D., Udvardi, M.K., Raffaele, S., Mongrand, S., Cullimore, J., Gamas, P., Niebel, A., and Ott, T. (2010). A remorin protein interacts with symbiotic receptors and regulates bacterial infection. Proceedings of the National Academy of Sciences of the United States of America 107, 2343–2348.

Levy, J., Bres, C., Geurts, R., Chalhoub, B., Kulikova, O., Duc, G., Journet, E.P., Ane, J.M., Lauber, E., Bisseling, T., Denarie, J., Rosenberg, C., and Debelle, F. (2004). A putative Ca^2+^ and calmodulin-dependent protein kinase required for bacterial and fungal symbioses. Science 303, 1361–1364.

Libault, M., Farmer, A., Brechenmacher, L., Drnevich, J., Langley, R.J., Bilgin, D.D., Radwan, O., Neece, D.J., Clough, S.J., May, G.D., and Stacey, G. (2010). Complete Transcriptome of the Soybean Root Hair Cell, a Single-Cell Model, and Its Alteration in Response to Bradyrhizobium japonicum Infection. Plant Physiology 152, 541–552.

Limpens, E., Moling, S., Hooiveld, G., Pereira, P.A., Bisseling, T., Becker, J.D., and Küster, H. (2013). Cell- and Tissue-Specific Transcriptome Analyses of Medicago truncatula Root Nodules. PLoS ONE 8, e64377.

Lin, J., Frank, M., and Reid, D. (2020). No Home without Hormones: How Plant Hormones Control Legume Nodule Organogenesis. Plant Communications 1.

Liu, C.-W., Breakspear, A., Stacey, N., Findlay, K., Nakashima, J., Ramakrishnan, K., Liu, M., Xie, F., Endre, G., and de Carvalho-Niebel, F. (2019a). A protein complex required for polar growth of rhizobial infection threads. Nature communications 10, 2848.

Liu, C.-W., Breakspear, A., Guan, D., Cerri, M.R., Jackson, K., Jiang, S., Robson, F., Radhakrishnan, G.V., Roy, S., Bone, C., Stacey, N., Rogers, C., Trick, M., Niebel, A., Oldroyd, G.E.D., De Carvalho-Niebel, F., and Murray, J.D. (2019b). NIN Acts as a Network Hub Controlling a Growth Module Required for Rhizobial Infection. Plant Physiology 179, 1704–1722.

Liu, H., Zhang, C., Yang, J., Yu, N., and Wang, E. (2018). Hormone modulation of legume-rhizobial symbiosis. Journal of Integrative Plant Biology 60, 632–648.

Liu, M., Kameoka, H., Oda, A., Maeda, T., Goto, T., Yano, K., Soyano, T., and Kawaguchi, M. (2022). The effects of ERN1 on gene expression during early rhizobial infection in Lotus japonicus. Frontiers in Plant Science 13.

Love, M.I., Huber, W., and Anders, S. (2014) Moderated estimation of fold change and dispersion for RNA-seq data with DESeq2, Genome biology 15 (2), 1–21.

Mathesius, U., Schlaman, H.R., Spaink, H., Sautter, C., Rolfe, B., and Djordjevic, M.A. (1998). Auxin transport inhibition precedes root nodule formation in white clover roots and is regulated by flavanoids and derivatives of chitin oligosaccharides. The Plant Journal 14, 23–34.

Mergaert, P., Kereszt, A., and Kondorosi, E. (2020). Gene Expression in Nitrogen-Fixing Symbiotic Nodule Cells in Medicago truncatula and Other Nodulating Plants. The Plant Cell 32, 42–68.

Mortier, V., Den Herder, G., Whitford, R., Van de Velde, W., Rombauts, S., D’Haeseleer, K., Holsters, M., and Goormachtig, S. (2010). CLE peptides control Medicago truncatula nodulation locally and systemically. Plant Physiol 153, 222–237.

Murray, J.D. (2016). The cell cycle in nodulation. Mol Cell Biol Growth Differ Plant Cells 13, 220–235.

Okamoto, S., Shinohara, H., Mori, T., Matsubayashi, Y., and Kawaguchi, M. (2013). Root-derived CLE glycopeptides control nodulation by direct binding to HAR1 receptor kinase. Nat Commun 4, 2191.

Oldroyd, G.E. (2013). Speak, friend, and enter: signalling systems that promote beneficial symbiotic associations in plants. Nat Rev Microbiol 11, 252–263.

Oldroyd, G.E., and Long, S.R. (2003). Identification and characterization of nodulation-signaling pathway 2, a gene of Medicago truncatula involved in Nod actor signaling. Plant Physiol 131, 1027–1032.

Peck, S.C., Olson, D.C., and Kende, H. (1993). A cDNA sequence encoding 1-aminocyclopropane-1-carboxylate oxidase from pea. Plant Physiology 101, 689.

Penmetsa, R.V., and Cook, D.R. (1997). A Legume Ethylene-Insensitive Mutant Hyperinfected by Its Rhizobial Symbiont. Science 275, 527–530.

Penmetsa, R.V., Frugoli, J., Smith, L., Long, S.R., and Cook, D. (2003). Genetic evidence for dual pathway control of nodule number in Medicago truncatula. Plant Physiology 131, 998–1008.

Penmetsa, R.V., Uribe, P., Anderson, J., Lichtenzveig, J., Gish, J.C., Nam, Y.W., Engstrom, E., Xu, K., Sckisel, G., Pereira, M., Baek, J.M., Lopez-Meyer, M., Long, S.R., Harrison, M.J., Singh, K.B., Kiss, G.B., and Cook, D.R. (2008). The Medicago truncatula ortholog of Arabidopsis EIN2, sickle, is a negative regulator of symbiotic and pathogenic microbial associations. Plant Journal 55, 580–595.

Pislariu, C.I., Sinharoy, S., Torres-Jerez, I., Nakashima, J., Blancaflor, E.B., and Udvardi, M.K. (2019). The nodule-specific PLAT domain protein NPD1 is required for nitrogen-fixing symbiosis. Plant physiology 180, 1480–1497.

Roux, B., Rodde, N., Jardinaud, M.F., Timmers, T., Sauviac, L., Cottret, L., Carrère, S., Sallet, E., Courcelle, E., and Moreau, S. (2014). An integrated analysis of plant and bacterial gene expression in symbiotic root nodules using laser-capture microdissection coupled to RNA sequencing. The Plant Journal 77, 817–837.

Roy, S., and Müller, L.M. (2022). A rulebook for peptide control of legume–microbe endosymbioses. Trends in plant science.

Roy, S., Liu, W., Nandety, R.S., Crook, A., Mysore, K.S., Pislariu, C.I., Frugoli, J., Dickstein, R., and Udvardi, M.K. (2020). Celebrating 20 years of genetic discoveries in legume nodulation and symbiotic nitrogen fixation. The Plant Cell 32, 15–41.

Schiessl, K., Lee, T., Orvosova, M., Bueno-Batista, M., Stuer, N., Bailey, P.C., Wen, J., Mysore, K., and Oldroyd, G.E. (2023). Light sensitive short hypocotyl (LSH) confer symbiotic nodule identity in the legume Medicago truncatula. bioRxiv, 2023.2002. 2012.528179.

Schiessl, K., Lilley, J.L.S., Lee, T., Tamvakis, I., Kohlen, W., Bailey, P.C., Thomas, A., Luptak, J., Ramakrishnan, K., Carpenter, M.D., Mysore, K.S., Wen, J., Ahnert, S., Grieneisen, V.A., and Oldroyd, G.E.D. (2019). NODULE INCEPTION Recruits the Lateral Root Developmental Program for Symbiotic Nodule Organogenesis in Medicago truncatula. Current Biology 29, 3657–3668.e3655.

Schnabel, E., Chavan, S., Poehlman, W., Feltus, F.A., and Frugoli, J. (2023). Transcriptome analysis of Medicago truncatula Autoregulation of Nodulation mutants reveals that disruption of the SUNN pathway causes constitutive expression changes in a small group of genes, but the overall response to rhizobia resembles wild type, including induction of TML1 and TML2. bioRxiv, 2023.2001. 2019.524769.

Shinohara, H., and Matsubayashi, Y. (2010). Arabinosylated glycopeptide hormones: new insights into CLAVATA3 structure. Current Opinion in Plant Biology 13, 515–519.

Soyano, T., Liu, M., Kawaguchi, M., and Hayashi, M. (2021). Leguminous nodule symbiosis involves recruitment of factors contributing to lateral root development. Current Opinion in Plant Biology 59, 102000.

Suzaki, T., Yano, K., Ito, M., Umehara, Y., Suganuma, N., and Kawaguchi, M. (2012). Positive and negative regulation of cortical cell division during root nodule development in Lotus japonicus is accompanied by auxin response. Development 139, 3997–4006.

Tian, Y., Liu, W., Cai, J., Zhang, L.-Y., Wong, K.-B., Feddermann, N., Boller, T., Xie, Z.-P., and Staehelin, C. (2013). The nodulation factor hydrolase of Medicago truncatula: characterization of an enzyme specifically cleaving rhizobial nodulation signals. Plant physiology 163, 1179–1190.

Trujillo, D.I., Silverstein, K.A., and Young, N.D. (2019). Nodule-specific PLAT domain proteins are expanded in the Medicago lineage and required for nodulation. New Phytologist 222, 1538–1550.

van Noorden, G.E., Ross, J.J., Reid, J.B., Rolfe, B.J., and Mathesius, U. (2006). Defective long-distance auxin transport regulation in the Medicago truncatula super numeric nodules mutant. Plant Physiology 140, 1494–1506.

Venkateshwaran, M., Jayaraman, D., Chabaud, M., Genre, A., Balloon, A.J., Maeda, J., Forshey, K., den Os, D., Kwiecien, N.W., Coon, J.J., Barker, D., and Ane, J.M. (2015). A role for the mevalonate pathway in early symbiotic signaling. Proceedings of the National Academy of Science of the United States of America 112, 9781–9786.

Vernie, T., Kim, J., Frances, L., Ding, Y., Sun, J., Guan, D., Niebel, A., Gifford, M.L., de Carvalho-Niebel, F., and Oldroyd, G.E. (2015). The NIN Transcription Factor Coordinates Diverse Nodulation Programs in Different Tissues of the Medicago truncatula Root. Plant Cell 27, 3410–3424.

Waese, J., Fan, J., Pasha, A., Yu, H., Fucile, G., Shi, R., Cumming, M., Kelley, L.A., Sternberg, M.J., and Krishnakumar, V. (2017). ePlant: visualizing and exploring multiple levels of data for hypothesis generation in plant biology. The Plant Cell 29, 1806–1821.

Waese-Perlman, B., Pasha, A., Ho, C., Azhieh, A., Liu, Y., Sullivan, A., Lau, V., Esteban, E., Waese, J., Ly, G., Hooper, C., Staton, S.E., Brereton, N., Le, C., Nelson, R., Lumba, S., Goodstein, D., Millar, A.H., Parkin, I., Lukens, L., Ehlting, J., Rieseberg, L., Pitre, F., Brown, A., and Provart, N.J. (2021). ePlant in 2021: New Species, Viewers, Data Sets, and Widgets.

Yang, J., Lan, L., Jin, Y., Yu, N., Wang, D., and Wang, E. (2022). Mechanisms underlying legume–rhizobium symbioses. Journal of Integrative Plant Biology 64, 244–267.

