## Supplemental Figure 1 for "A laser capture microdissection transcriptome of *M. truncatula* roots responding to rhizobia reveals spatiotemporal tissue expression patterns of genes involved in nodule signaling and organogenesis"

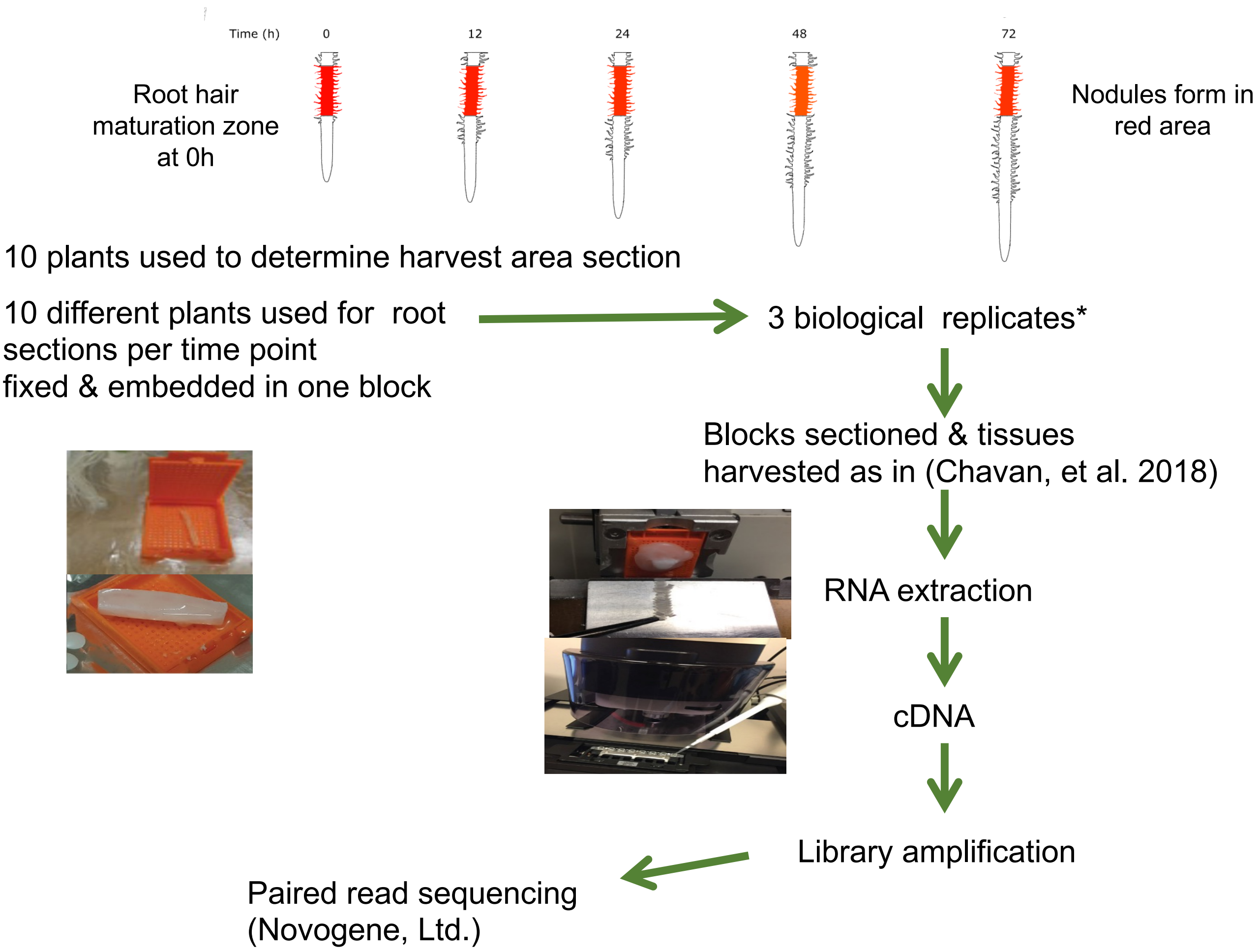


**Supplemental Figure 1. Diagram of procedure designed to increase signal to noise ratio** Twenty plants per genotype (t = 0 hpi) were collected and rhizobia added to the growth apparatus immediately after collection of the 0 hpi samples. Additional samples of 20 plants per genotype were collected at 12, 24, 48, and 72 hpi. Ten plants from each collection were used to determine average root length and 2 cm segments representing the zone of development of the first nodules were collected from the remaining 10 plants. At 0 hpi this region started 1 cm from the root tip, where the first full length root hairs were present. At later time points, this region was determined by calculating the average root growth since $t$=0 and adding this distance to 1 cm.
