## Supplemental Figure 2 for "A laser capture microdissection transcriptome of *M. truncatula* roots responding to rhizobia reveals spatiotemporal tissue expression patterns of genes involved in nodule signaling and organogenesis"

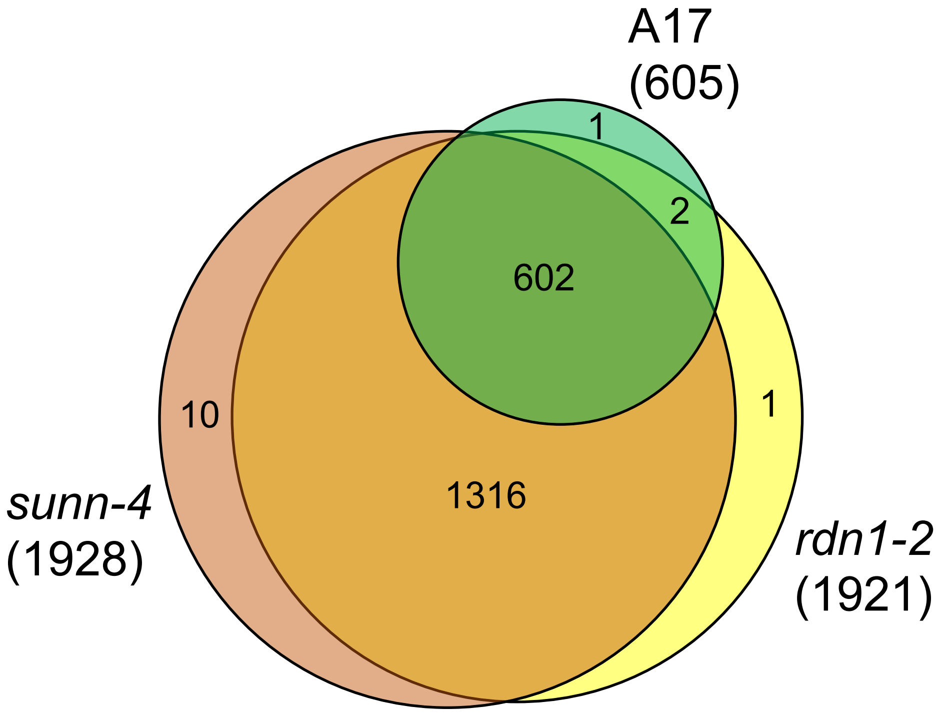


**Supplemental Figure 2. Overlap of rhizobial response genes between wild type and supernodulation mutants.** 1932 genes were identified as increasing in response to rhizobia in wild type (A17), *sunn-4*, or *rdn1-2* during the 72 hpi (from data in Schnabel et al., 2023). The Venn diagram shows the high degree of similarity between genes responding in the three lines. The number of genes unique to any line or in common between two or lines is written within the space.
