## Supplemental Figure 3 for "A laser capture microdissection transcriptome of *M. truncatula* roots responding to rhizobia reveals spatiotemporal tissue expression patterns of genes involved in nodule signaling and organogenesis"

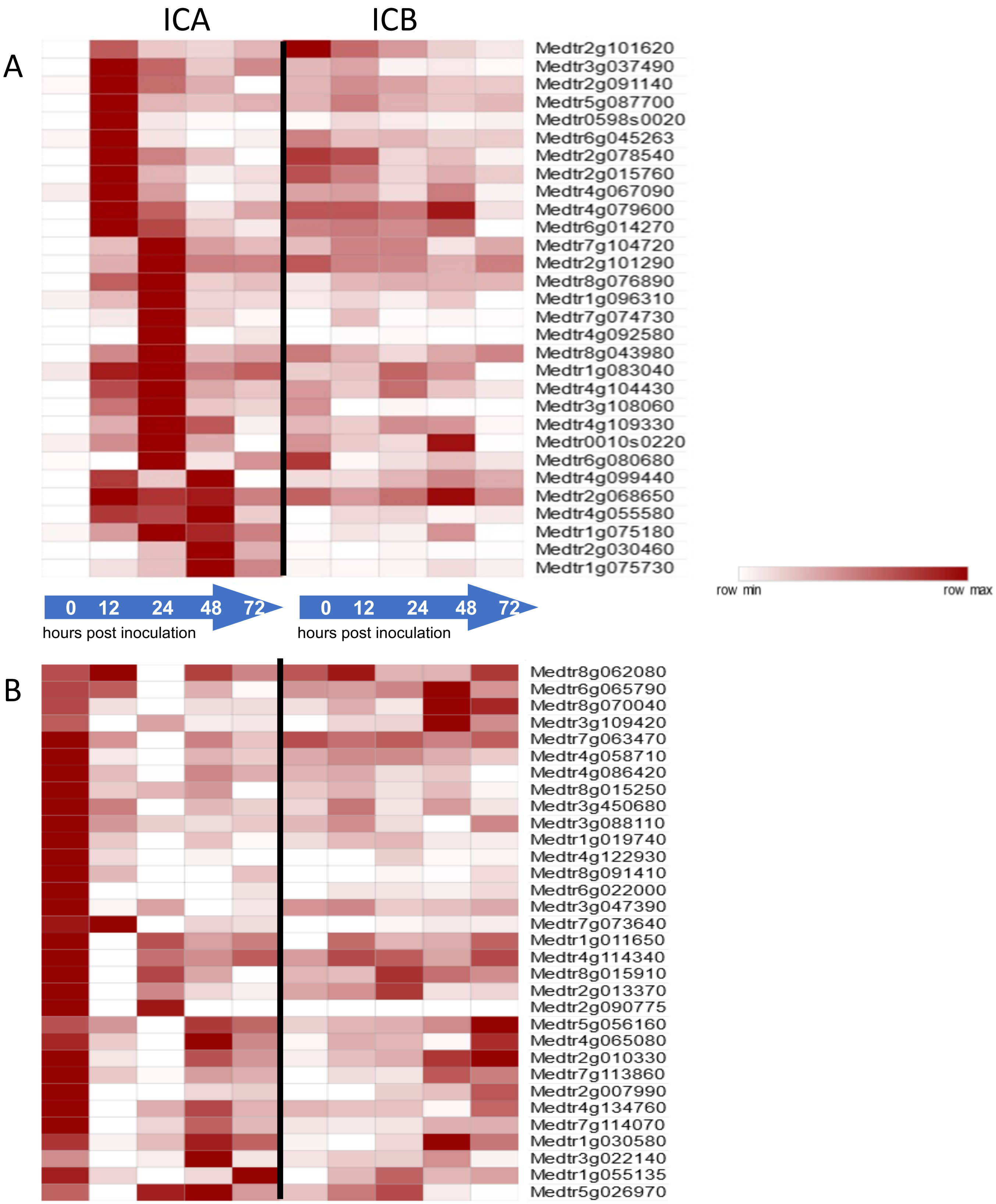


**Supplemental Figure 3. Genes with differential expression in spatially differentiated inner cortical cells at early timepoints**. (A) Genes induced early (12 or 24 hpi) in the ICA but not the ICB. (B) Genes induced early (12 or 24 hpi) in the ICB but not the ICA. Annotations of genes occur in the searchable lists in Supplemental Dataset 2D.
