## Supplemental Figure 4 for "A laser capture microdissection transcriptome of *M. truncatula* roots responding to rhizobia reveals spatiotemporal tissue expression patterns of genes involved in nodule signaling and organogenesis"

| | |
|---|---|
| Medtr2g011040 | viral movement protein |
| Medtr7g099030 | DENN (AEX-3) domain protein |
| Medtr8g062630 | transmembrane protein, putative |
| Medtr6g009090 | ABC transporter transmembrane region protein |
| Medtr2g071960 | Rpp4C4 |
| Medtr3g077110 | myb transcription factor |
| Medtr4g062460 | disease resistance response/dirigent-like protein |
| Medtr8g021350 | receptor-like kinase, putative |
| Medtr3g005495 | B3 DNA-binding domain protein |

| | |
|---|---|
| Medtr8g028500 | Rpp4C1 |
| Medtr5g007740 | chalcone and stilbene synthase family protein |
| Medtr3g101720 | PIF1 helicase, putative |
| Medtr6g045353 | hypothetical protein |

| | |
|---|---|
| Medtr1g029070 | nuclear transcription factor Y protein |
| Medtr6g074780 | disease resistance protein (TIR-NBS-LRR class), putative |

| | |
|---|---|
| Medtr8g010090 | polygalacturonase/glycoside hydrolase family protein |
| Medtr2g019610 | cysteine protease, putative |
| Medtr4g072140 | hypothetical protein |
| Medtr4g023510 | PPR containing plant-like protein |
| Medtr1g111010 | Pmr5/Cas1p GDSL/SGNH-like acyl-esterase family protein |
| Medtr3g069345 | hypothetical protein |

| | |
|---|---|
| Medtr5g070980 | cytochrome P450 family 71 protein |
| Medtr3g073660 | hypothetical protein |
| Medtr0336s0070 | hypothetical protein |
| Medtr4g019350 | transmembrane protein, putative |

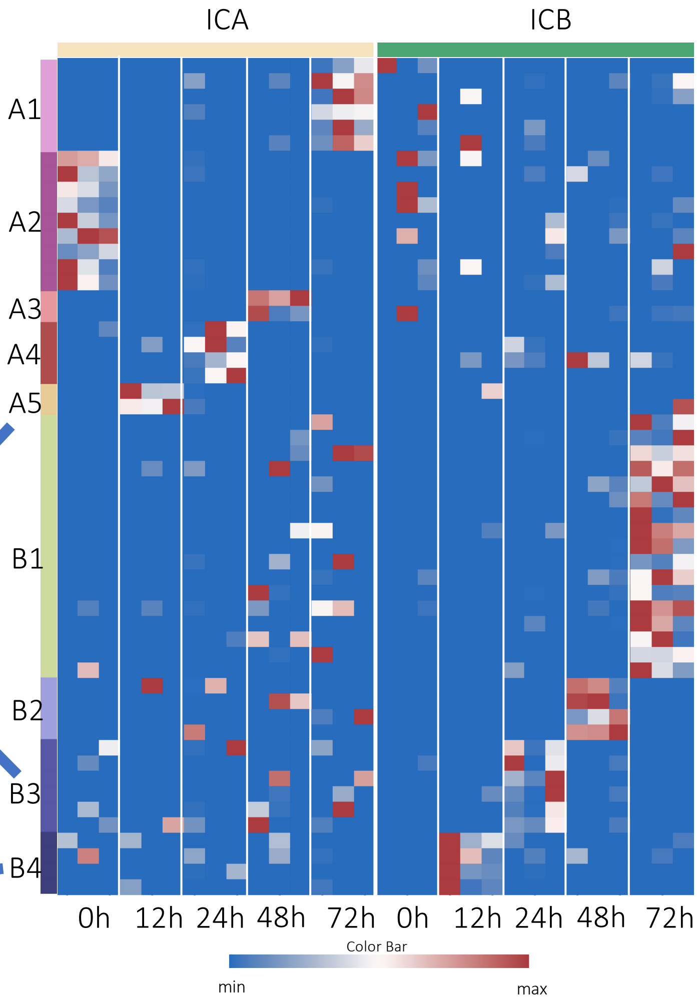


**Supplemental Figure 4. Gene expression profiles in ICA and ICB in inoculated *M. truncatula* over 72 hpi.** (A) Heatmap of 52 genes from Supplemental Dataset 2E with different temporal expression patterns between ICA and ICB as identified by GeneShift. Data are presented using min-max scaled log2 (x+1) transformed FPKM expression values. Each row represents a gene, and each column represents the expression profile for a single biological sample, with 3 columns for each gene/timepoint. GeneShift trajectory groups (displayed in Supplemental Figure 4) are numbered on the Y axis with hours post inoculation (hpi) on the X axis. The color scale reflects relative gene expression within a row. Genes from trajectory lists discussed in text are indicated with arrows.
