## Supplemental Figure 5 for "A laser capture microdissection transcriptome of *M. truncatula* roots responding to rhizobia reveals spatiotemporal tissue expression patterns of genes involved in nodule signaling and organogenesis"

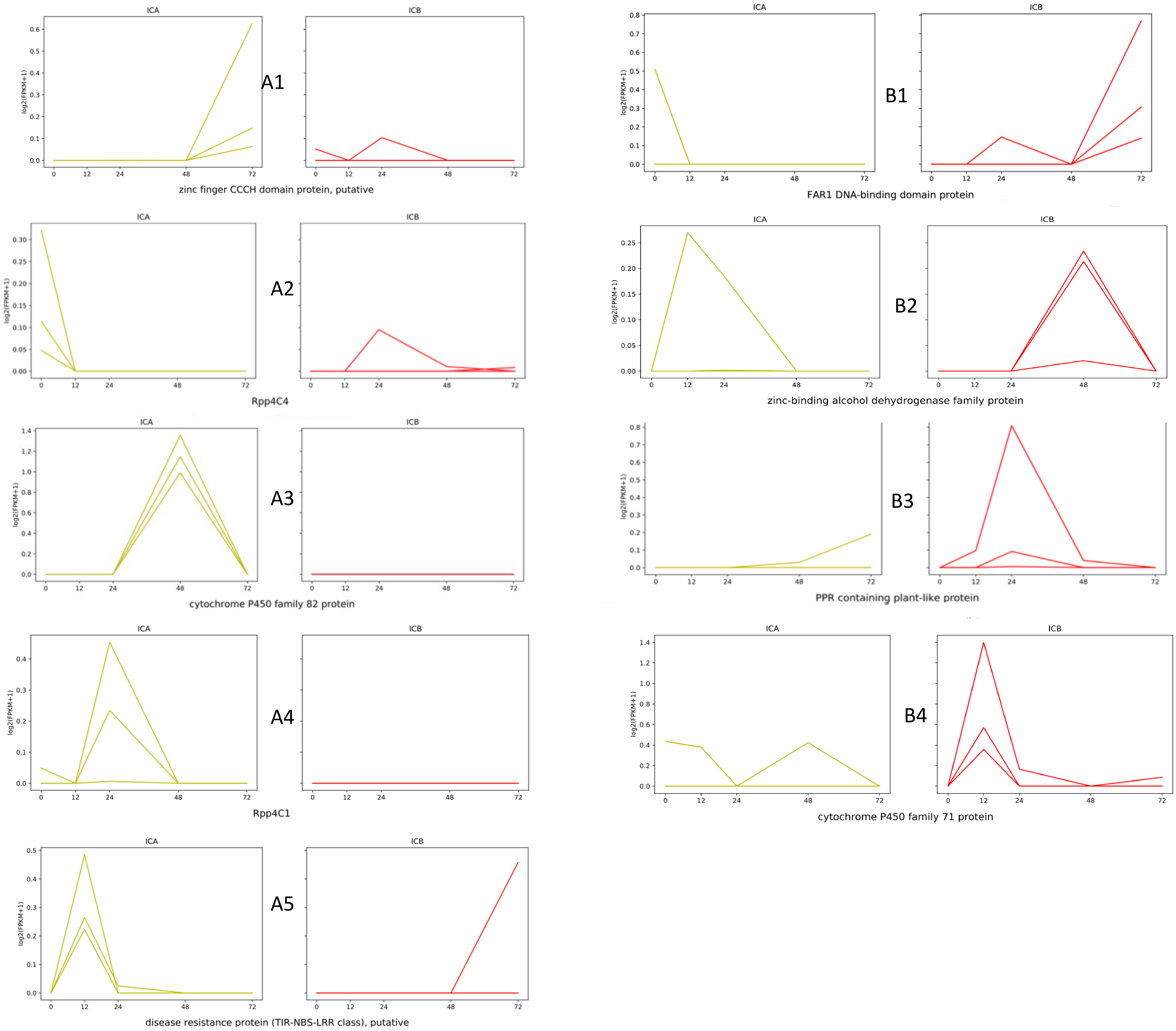


**Supplemental Figure 5. Expression profiles of representative genes in GeneShift trajectory sets for ICA and ICB tissues.** Example gene expression data for each trajectory set in the text. “A” sets have dynamic expression in ICA cells (yellow) versus ICB cells (red), while “B” sets have dynamic expression in ICB cells versus ICA cells. Data are presented using min-max scaled log2 (x+1) transformed FPKM expression values (Y axis) and time post inoculation (X axis). Each line indicates expression in a single replicate; there are three lines in each plot but some overlap. The color scale reflects relative gene expression within a row.
